# Integrative phenotypic and transcriptomic validation of an alveolar-like macrophage model reveals early host–pathogen dynamics during *Aspergillus fumigatus* infection

**DOI:** 10.64898/2026.04.30.721838

**Authors:** Jeany Söhnlein, Sascha Schäuble, Juan Prada Salcedo, Zahraa Abboud, Dalia Sheta, Kerstin Hünniger-Ast, Michelle Seif, Hermann Einsele, Thomas Dandekar, Gianni Panagiotou, Andreas Beilhack, Juergen Loeffler

## Abstract

Primary alveolar macrophages (pAMs) are essential for the rapid clearance of conidia and maintenance of pulmonary homeostasis. However, *Aspergillus fumigatus* remains the leading cause of invasive pulmonary aspergillosis in immunocompromised patients, and the mechanisms governing fungal clearance versus invasion remain poorly understood. Although, pAMs can be isolated from human donors, their broader application in *in vitro* infection studies is limited by their low availability and technical challenges with their maintenance in culture.

In this study, we successfully adapted a previously established monocyte-derived alveolar-like macrophage (ALM) model to investigate early host-pathogen interactions upon *A. fumigatus* challenge. Given the requirement of GM-CSF for maintaining alveolar macrophage identity and function, we included GM-CSF differentiated macrophages (GM-M), as a widely employed reference model. Primary alveolar macrophages (pAM), isolated from human lung biopsies were utilized to validate the physiological relevance of the ALM model. Combined phenotypic, transcriptomic and functional analyses demonstrated that ALMs closely resemble pAMs under both steady-state and infection conditions across multiple time points and fungal burdens. Notably, fungal dual RNA-sequencing revealed a significant upregulation of fungal virulence-associated factors during interaction with ALM, which was not observed in our GM-M co-cultures.

Collectively, these findings support the use of ALMs as a robust, experimentally accessible and physiologically relevant *in vitro* model for investigating early *A. fumigatus* infection, providing new insights into host-pathogen dynamics at the alveolar interface.

## 1 Introduction

*Aspergillus fumigatus,* a saprotrophic mold commonly found in soil and decaying organic matter, produces 2-3 µm small airborne conidia. Its thermotolerance, broad pH adaptability, and hydrophobic conidial surface contribute to its success as an opportunistic airborne pathogen. Upon inhalation, conidia can reach the lung alveoli, where they embed within the surfactant layer of the alveolar epithelium and initiate germination within hours (Ochs et al. 2020; van de Veerdonk et al. 2017). In immunocompromised individuals, this can lead to invasive pulmonary aspergillosis (IPA), a life-threatening disease particularly dangerous for patients with neutropenia, undergoing chemotherapy or transplantation, or receiving prolonged corticosteroid therapy (Corre et al. 2010; Elkhapery, Fatima, and Soubani 2025).

Airway epithelial cells and primary alveolar macrophages (pAMs) constitute the first line of defense against *A. fumigatus*. These cells employ multiple antifungal mechanisms, including conidial phagocytosis, production of cytokines and antimicrobial peptides, and generation of reactive oxygen species (Osherov 2012). As the dominant immune cell population in the alveolar niche under homeostatic conditions, pAMs act as key regulators of pulmonary immunity, with their phenotype and function shaped by the local tissue environment. Upon infection, they coordinate early immune responses by promoting leukocyte recruitment and directly contributing to pathogen clearance (Gosselin et al. 2014; Malainou et al. 2023; Pahari et al. 2023).

Although pAMs play a central role in the early immune response against *A. fumigatus* (Margalit and Kavanagh 2015), underlying mechanisms remain incompletely understood. Experimental approaches have included *in vitro* and *in vivo* infection models as well as the use of isolated pAMs (Rosowski et al. 2018; Bhatia et al. 2011; Wang et al. 2022; Amich et al. 2020; Kalleda et al. 2016; Hellmann et al. 2017). However, these strategies face major limitations. pAMs rapidly lose their tissue-specific phenotype during *in vitro* culture, and their isolation is constrained by low yield, donor variability, and limited amenability to genetic manipulation (Pahari et al. 2023; Papp et al. 2018; Malainou et al. 2023). To address these challenges, recent work has established a monocyte-derived alveolar-like macrophage model (ALM) that recapitulates key features of pAMs while offering improved scalability and experimental accessibility. Although the ALM model has previously been validated in bacterial and viral infection settings, its applicability to fungal pathogens has not been investigated (Pahari et al. 2023; Pahari et al. 2024).

Here, we adapted this previously described ALM model to study host-pathogen interactions during *A. fumigatus* infection and systematically compared it to biopsy isolated human pAMs and GM-CSF differentiated macrophages (GM-Ms) derived from the same donor as ALMs. GM-Ms were included as a widely used reference model, as GM-CSF is critical for pAM development and function in the lung environment (Malainou et al. 2023; Pahari et al. 2023). Using combined phenotypic, transcriptomic and functional analyses, we demonstrate that ALMs closely resemble pAMs both under steady-state and infectious conditions, independent of fungal burden and infection time. Hence, we demonstrate that ALMs are suitable to investigate early host-pathogen interactions in *A. fumigatus* infection.

## 2 Materials and Methods

### Cell culture

CD14^+^ monocytes were isolated from peripheral blood mononuclear cells (PBMCs) as previously described (Seelbinder et al. 2020; Heilig et al. 2025). Briefly, PBMCs were obtained from healthy adults using leukoreduction chambers. Mononuclear cells were isolated by density-gradient centrifugation using Histopaque (density 1.077 g/mL; Sigma) and subsequently quantified using a Vi-Cell XR counter (Beckman Coulter). Employing a positive selection with human CD14 microbeads (Miltenyi Biotec) from PBMCs, CD14^+^ monocytes were isolated, according to manufacturer’s protocol. Differentiation into GM-Ms was achieved by supplementing Macrophage Serum-free medium (M-SFM) (Gibco) with three doses of 20 ng/mL GM-CSF (Miltenyi), while ALMs were generated by supplementing M-SFM with three doses of 100 µg/mL Curosurf (Chiesi), 10 ng/mL GM-CSF, 5 ng/mL IL-10 (Miltenyi) and 5 ng/mL TGF-β (Miltenyi) on day 0, 2, and 4. Macrophages were harvested on day 6 with accutase (Sigma) for 20 min at 37°C, 5% CO_2_ and a cell scraper and used for further experiments. pAMs were isolated from fresh human lung biopsies provided by the University Hospital Wuerzburg. Concisely, lung tissue was dissected with scissors and pAMs were washed out with PBS (Sigma) and filtered through a 70 µm strainer. Lysis of erythrocytes was achieved with erythrocyte lysis buffer (QIAGEN) for 5 min at room temperature (RT). Cells were washed once with RPMI 1640 (Gibco) + 10% FBS + 120 µg/mL gentamicin, resuspended and seeded overnight in a 6-well plate. Afterwards, adherent cells were washed three times with PBS before harvesting with accutase and a cell scraper, centrifuged and resuspended in RPMI 1640 + 10% FBS + 120 µg/mL gentamicin for subsequent experiments. Lung samples from heavy smokers were excluded. Use and processing of human blood specimen (#302/12) and pAMs (#34/22) from tissue was approved by the Ethics Committee of the University Hospital Wuerzburg and informed consent of lung biopsy donors was obtained from each patient before surgery.

### Aspergillus fumigatus culture conditions

*A. fumigatus* ATCC46645, CEA17 Δ*akuB KU80*, Δ*aspf1*, and Δ*aspf2* were cultured on malt agar plates as previously described (Krespach et al. 2023) with or without supplementation of 0.1 µg/mL pyrithiamine (Sigma) at 37°C for 4 days. *A. fumigatus* DsRed^+^ or Af293 were grown on *Aspergillus* minimal media (Scott et al. 1982; Brakhage and Van den Brulle 1995) and incubated at 37°C for 7 days. The medium was supplemented with 100 µg/mL hygromycin (Sigma) for DsRed^+^ conidia. Fungal conidia were harvested using sterile water and filtered through a 40 µm cell strainer. Conidia were kept at 4°C and used for a maximum of 2 weeks.

To assess the influence of Asp f1 and Asp f2 deficiency on the germination of *A. fumigatus,* 1x10^5^ conidia of *A. fumigatus* CEA17, *Δaspf1* and *Δaspf2* were inoculated in RPMI 1640 and incubated at 37°C, 5% CO_2_ for 4h-9h. At each time point, 10 pictures were taken using a Zeiss Primovert Microscope using a 40x objective lens. During data acquisition, clumps of conidia/hyphae were not considered (Jia et al. 2025).

### Phenotypic evaluation of pAM, ALM and GM-M

For immunofluorescence analysis, pAM, ALM or GM-M were blocked with 5% BSA/PBS for 10 min at 4°C and stained with anti-CD206 AF488 (1:20; Biolegend) and anti-CD11c AF647 (1:10; Biolegend) at 4°C for 20 min. Cells were washed three times with PBS, resuspended in RPMI 1640 + 10% FBS + 120 µg/mL gentamicin and 1x10^5^ macrophages were seeded into µ-Slide VI 0.4 chambers (ibidi). After overnight incubation at 37°C, 5% CO_2_, cells were stained with Hoechst (1:2,000; Thermo Fisher) in PBS for 10 min at RT followed by three times washing with PBS. Supernatant was removed and exchanged to RPMI 1640 + 10% FBS + 120 µg/mL gentamicin and imaged using a Zeiss LSM780 confocal microscope (Zeiss, Germany).

For flow cytometry, 1x10^5^ cells were blocked with human Fc receptor blocking reagent (Miltenyi), according to manufacturer’s protocol, at 4°C for 10 min and stained with fixable Viability Dye eFluor^TM^ 780 (1:2,000; Invitrogen^TM^), anti-HLA-DR-DP-DQ VioB515 (1:100; Miltenyi), anti-CD206 PECy7 (1:100; Biolegend), anti-CD11b APC (1:50; Miltenyi) or the respective isotype control at 4°C for 20 min. Inside staining was performed using the Inside Stain kit (Miltenyi) according to manufacturer’s protocol and anti-CD68 BV785 (1:20; Biolegend). Samples were analyzed on a CytoFlex Flow Cytometer (Beckman Coulter) and data was processed using Kaluza V2.2.1 software (Beckman Coulter).

### Macrophage infection assays

pAM, ALM and GM-M were infected with resting *A. fumigatus* ATC46645 conidia at a multiplicity of infection (MOI) of 1 for 6 h. Alternatively, cells were infected at MOI 5 for 9 h using strains *A. fumigatus* ATCC46645 or CEA17, Δ*aspf1*, or Δ*aspf2*. For dual RNA-sequencing, ALM and GM-M were infected with *A. fumigatus* ATCC46645 at MOI 5 for 6 and 9h. Infection was synchronized by centrifugation at 200xg, 5 min before incubation. Following infection, cells were harvested by scraping and centrifugation at 300xg, 5 min. Supernatants or cell pellets in RNAprotect Cell Reagent (QIAGEN) were stored at -80°C until further processing.

### Phagocytosis and killing efficiency quantification

Surface labelling of DsRed^+^ conidia was conducted as previously described (Brunel et al. 2017; Jhingran et al. 2012). FLARE conidia were stored in PBS at 4°C for a maximum of 2 weeks. ALM and GM-M were infected with *A. fumigatus* FLARE at a MOI of 0.5, 1, or 3 or with Af293 MOI 3 for isotype control and centrifuged at 200xg, 5 min to initiate proper infection, followed by incubation for 2 or 6h at 37°C, 5% CO_2_. FLARE infected cells were counterstained with calcofluor white (CFW) in RPMI 1640 + 10% FBS + 120 µg/mL gentamicin (1:40; Sigma) in RPMI 1640 + 10% FBS + 120 µg/mL gentamicin for 10 min at 37°C, 5% CO_2_, to distinguish between extracellular and phagocytosed conidia. Macrophages were blocked with Fc receptor blocking solution and stained with fixable Viability Dye eFluor^TM^ 780 (1:2,000), anti-CD206 BV785 (1:20; Biolegend) and anti-CD11c PEV770 (1:50; Miltenyi). Phagocytosis and killing efficiency were analyzed on a CytoFlex Flow Cytometer and evaluated using Kaluza V2.2.1 software. For representative pictures, ALM were blocked with 5% BSA/PBS, stained with anti-CD206 AF488 (1:20; Biolegend) and seeded overnight into µ-Slide VI 0.4 chambers (ibidi). Cells were infected using FLARE conidia at MOI 3 and incubated for 6h before counterstaining with CFW. Microscopy pictures were obtained employing a Zeiss CLSM780 confocal microscope.

To quantify the phagocytosis of *A. fumigatus* CEA17, Δ*aspf1*, or Δ*aspf2* conidia, ALMs were infected with a MOI 5 of each fungal strain for 2h. Infected cells were treated as described above and phagocytic efficiency was quantified on a CytoFlex Flow Cytometer (Beckman Coulter) and evaluated using Kaluza V2.2.1 software.

### RNA isolation and dual bulk transcriptomic profiling

RNA was isolated using the RiboPure^TM^ RNA-purification kit yeast (Invitrogen) including DNAse treatment according to manufacturer’s protocol. Isolated RNA was measured on a NanoQuant Infinite M200 Pro plate reader (Tecan). RNA quality was checked using a 5200 Fragment Analyzer with the DNF-471-33-SS Total RNA 15nt kit (Agilent Technologies). DNAse-treated RNA was depleted of ribosomal RNA molecules using RiboCop for Human/Mouse/Rat V2 (Cat. No. 144) rRNA depletion kit (Lexogen). The ribo-depleted RNA samples were first fragmented using ultrasound (4 pulses of 30s at 4°C). Then, an oligonucleotide adapter was ligated to the 3’ end of the RNA molecules. First-strand cDNA synthesis was performed using M-MLV reverse transcriptase with the 3’ adapter as primer. After purification, the 5’ Illumina TruSeq sequencing adapter was ligated to the 3’ end of the antisense cDNA. The resulting cDNA was PCR-amplified using a high-fidelity DNA polymerase and the barcoded TruSeq-libraries were pooled in approximately equimolar amounts. The size distribution of the barcoded DNA libraries was estimated to be ∼400bp by electrophoresis on 5200 Fragment Analyzer with the DNF-474-33-HS NGS Fragment 1-6000bp kit (Agilent Technologies). Sequencing of pooled libraries, spiked with PhiX control library, was performed at 50 million reads per sample in single-ended mode with 100 cycles on the NovaSeq 6000 platform (Illumina). Demultiplexed FASTQ filed were generated with bcl-convert v4.2.4 (Illumina).

### Dual RNA-sequencing processing and pathway analysis

Raw sequencing reads including quality control and gene abundance estimation were processed using the GEO2RNAseq pipeline (v0.9.12) (Seelbinder et al. 2020) implemented in *R* (v3.5.1). Read quality was assessed before and after trimming using FastQC (v0.11.8). Low-quality bases and adapter sequences were removed with Trimmomatic (v0.36) and ribosomal RNA reads were filtered using SortMeRNA (v2.1) with a combined database containing all rRNA references provided by SortMeRNA. To enable simultaneous host-pathogen analysis, a combined reference genome was generated by merging the *Homo sapiens* reference genome (GRCh38, version 89) with the *A. fumigatus* Af293 reference genome (version s03-m05-r05) in FASTA format. Corresponding exon annotations were extracted and combined into a unified annotation file. The merged reference genome was indexed with exon information using HiSat2 (v2.1.0) in single-end mode) and gene abundance estimation was performed with featureCounts (v1.28.0) using default parameters. MultiQC (v1.7) was used to summarize the output of FastQC, Trimmomatic, HiSat, featureCounts and SAMtools. The read count matrix with gene abundances for both species without and with median-of-ratios normalization (MRN) (Anders and Huber 2010) were extracted. Differential gene expression per organism was analyzed using GEO2RNaseq. Pairwise tests were performed using four statistical tools (DESeq v1.30.0, DESeq2 v1.18.1, limma voom v3.34.6 and edgeR v3.20.7) to report p values and multiple testing corrected p values using the false-discovery rate method q = FDR(p) for each tool. Gene expression differences were considered significant if they were reported significant by all four tools (p ≤ 0.05) and a minimum fold change of log_2_(FC_MRN_) ≥ 1. Gene Ontology (GO) enrichment analysis was performed in *R* using the GSEGO function, with genes ranked according to log_2_ fold change values derived from DESeq2 differential expression analysis. For human gene annotation, the org.Hs.eg.db package from AnnotationDbi was used, while *A. fumigatus* Af293 annotations were obtained from a publicly available resource (Pardeshi 2021). Enriched GO terms were visualized using dot plots.

### Host-pathogen protein-protein interaction analysis

To prioritize biological processes relevant to host–pathogen interactions, enriched GO terms were filtered using a predefined curated list of interaction-associated terms provided in the Supplementary Table S1. This filtering included extracellular and cell wall–associated terms within the Cellular Component (CC) ontology as well as interaction-relevant Molecular Function (MF) categories. Differentially expressed genes (DEGs) were retained when annotated to selected GO terms in both CC and MF categories, ensuring consistency between subcellular localization and molecular function. Subsequently, we filtered for predicted protein features associated with host–pathogen interactions using established computational tools, including NetGPI-1.1 (GPI-anchor prediction) (Gíslason et al. 2021; Gíslason 2021), SignalP 6.0 (signal peptide prediction) (Teufel et al. 2022), TargetP 2.0 (secretory pathway targeting) (Almagro Armenteros et al. 2019), DeepTMHMM 1.0 (transmembrane domain prediction) (Hallgren et al. 2022), and DeepLoc 2.1 (subcellular localization and membrane protein classification) (Ødum et al. 2024). Proteins were retained if they contained at least one interaction-associated feature, including a GPI anchor, signal peptide, predicted secretion signal, appropriate subcellular localization, or one or more transmembrane domains. To further identify potential virulence-associated factors, *A. fumigatus* proteins were queried against the PHI-base database (Urban et al. 2025). *A. fumigatus* candidate genes were subsequently prioritized using a composite scoring approach integrating functional annotation and predicted protein features to enrich for proteins potentially involved in direct host–pathogen interactions. A pathogen relevance score was assigned to *A. fumigatus* proteins based on functional annotations and protein descriptions using a custom *R* function. Briefly, proteins were classified into five hierarchical tiers reflecting their potential involvement in host–pathogen interactions, with only the highest-ranking matching category retained per protein. Tier 1 (score = 10) included direct pathogenicity factors such as allergens, toxins, virulence-associated proteins, invasins, and adhesins, as well as annotations related to pathogenesis or immune evasion. Tier 2 (score = 8) comprised cell wall–degrading enzymes, including chitinases, glucanases, and non-ribosomal proteases/peptidases, along with corresponding GO terms. Tier 3 (score = 6) included enzymes involved in tissue degradation, such as cellulases, xylanases, and pectinases. Tier 4 (score = 4) captured proteins associated with host adaptation and stress responses, including catalases, superoxide dismutases, and heat shock proteins. Tier 5 (score = 3) included proteins predicted to be secreted or localized to the cell surface. Classification was performed by matching protein names and GO annotations (biological process, molecular function, and cellular component) using case-insensitive pattern recognition. Proteins not matching any category were assigned a score of 0 and labeled as not relevant. Finally, protein–protein interaction (PPI) networks were constructed using interaction data retrieved from the STRING database (DB) version 12.0. We used all possible interactions and a confidence threshold of 0.4.

### cDNA synthesis and quantitative reverse transcription PCR

To validate the fungal transcriptomic profile observed in the dual RNA-sequencing analysis, total RNA was isolated with the RiboPure^TM^ RNA-purification kit yeast (Invitrogen) including DNAse treatment according to manufacturer’s protocol. The RNA was converted into cDNA using the High-Capacity cDNA Reverse Transcription Kit (Applied Biosystems^TM^). qRT-PCR reactions were performed in duplicates employing 10 nM gene-specific primers and iTaq^TM^ Universal SYBR® Green Supermix (BioRad Laboratories) on a StepOnePlus^TM^ Real-Time PCR System (Applied Biosystems^TM^). Relative gene expression was calculated using the ΔΔCt method relative to the house-keeping gene *act1*. Primers are listed in the supplementary table S2. Primers for *act1*, *gliZ*, *cat1*, and *cipC* were selected from published data (Canela, Takami, and da Silva Ferreira 2017; Seelbinder et al. 2020; Sueiro-Olivares et al. 2015).

### Cytokine analysis

Custom ProcartaPlex (Thermo Fisher) or custom LEGENDplex (Biolegend) assay kits were used for multiplexed quantification of cytokine concentrations in culture supernatants according to the manufacturer’s protocol. ProcartaPlex data acquisition and analysis were performed using a Bio-Plex 200 Luminex reader in combination with Bio-Plex Manager Software version 6.2 (Bio-Rad). LEGENDplex data acquisition and analysis was performed on a CytoFlex Flow Cytometer (Beckman Coulter) according to manufacturer’s protocol. CCL2 concentrations measured out-of-range high were capped at the top standard of 10,000 pg/mL. Furthermore, due to the high occurrence of IL-8 concentrations, which were out-of-range high, a single-plex ELISA was performed for pAMs, ALMs and GM-Ms infected with *A. fumigatus* CEA17, *Δaspf1* and *Δaspf2* MOI 5 for 9h. Single-plex ELISA assays including ELISA MAX Deluxe Set Human for TNF, IL-8, CXCL10 (Biolegend) or Human DuoSet kits for CCL3 and CCL4 (R&D Systems) were performed according to manufacturer’s protocol.

### Colony forming unit analysis

ALMs were infected with MOI 5 of *A. fumigatus* CEA17, *Δaspf1* and *Δaspf2* for 9h. Subsequently, cells were lysed with cold ddH_2_O. The conidia suspensions were quantitatively cultured by serial dilution, plated on malt agar plates and incubated at 37°C for 18h. Afterwards, the *A. fumigatus* fungal burdens (number of CFU/mL) were determined (Xiong et al. 2023). ALMs were chosen due to their pAM resembling cytokine profile during infection with each of the three tested strains.

### Statistical analysis

GraphPad Prism version 10, Kaluza version 2.2.1, and *R* version 3.5.1 were used for data compilation, analysis and visualization. Depending on the data format, significance testing was performed using Kruskal-Wallis Test with Dunn’s multiple comparison, Ordinary One-Way ANOVA with Sidák’s or Tukey’s Multiple Comparison, RM-ANOVA with Dunnet’s Multiple Comparison, or Two-Way ANOVA with Tukey’s Multiple Comparison. The specific significance tests are specified in the figure legends. P-Values < 0.05 were considered significant.

## 3 Results

### 3.1 Alveolar-like macrophages resemble primary alveolar macrophages under steady-state and *A. fumigatus* infection conditions

To investigate early alveolar macrophage (Mφ) responses to *A. fumigatus*, we first assessed whether our differentiation protocol generated ALMs that exhibit key features of pAMs under steady-state and infection (Fig. 1A). Morphological analysis using immunofluorescence staining of established pAM markers (Fig. S1A) (Bain and MacDonald 2022) revealed that both pAMs and ALMs displayed a predominantly round morphology, consistent with previous reports (Pahari et al. 2023), whereas GM-Ms appeared flatter and more elongated (Fig. 1B). pAMs exhibited a larger cell size compared to ALMs, consistent with flow cytometry measurements (Fig. 1C). Moreover, pAMs showed higher autofluorescence, a characteristic feature of tissue-resident cells, whereas ALM and GM-M exhibited lower autofluorescence (Fig. 1D) likely reflecting their monocyte-derived origin (Knab et al. 2025). To further characterize Mφ identity, we analyzed the expression of canonical pAM markers, including signature markers CD206 and CD68 (Fig. S1B) (Sun, Cui, and Zhang 2026; Bain and MacDonald 2022). Flow cytometric analysis revealed gradual differences in marker expression across the three Mφ populations, particularly for HLA II and CD11b (Fig. 1D). Quantitative analysis of mean fluorescence identity (MFI) demonstrated that ALMs expressed HLA-DR-DP-DQ and CD68 at levels comparable to pAMs. For CD206 and CD11b, ALMs showed intermediate expression, more closely resembling pAMs than GM-Ms, which exhibit significant differences to pAMs.

**Figure 1:**
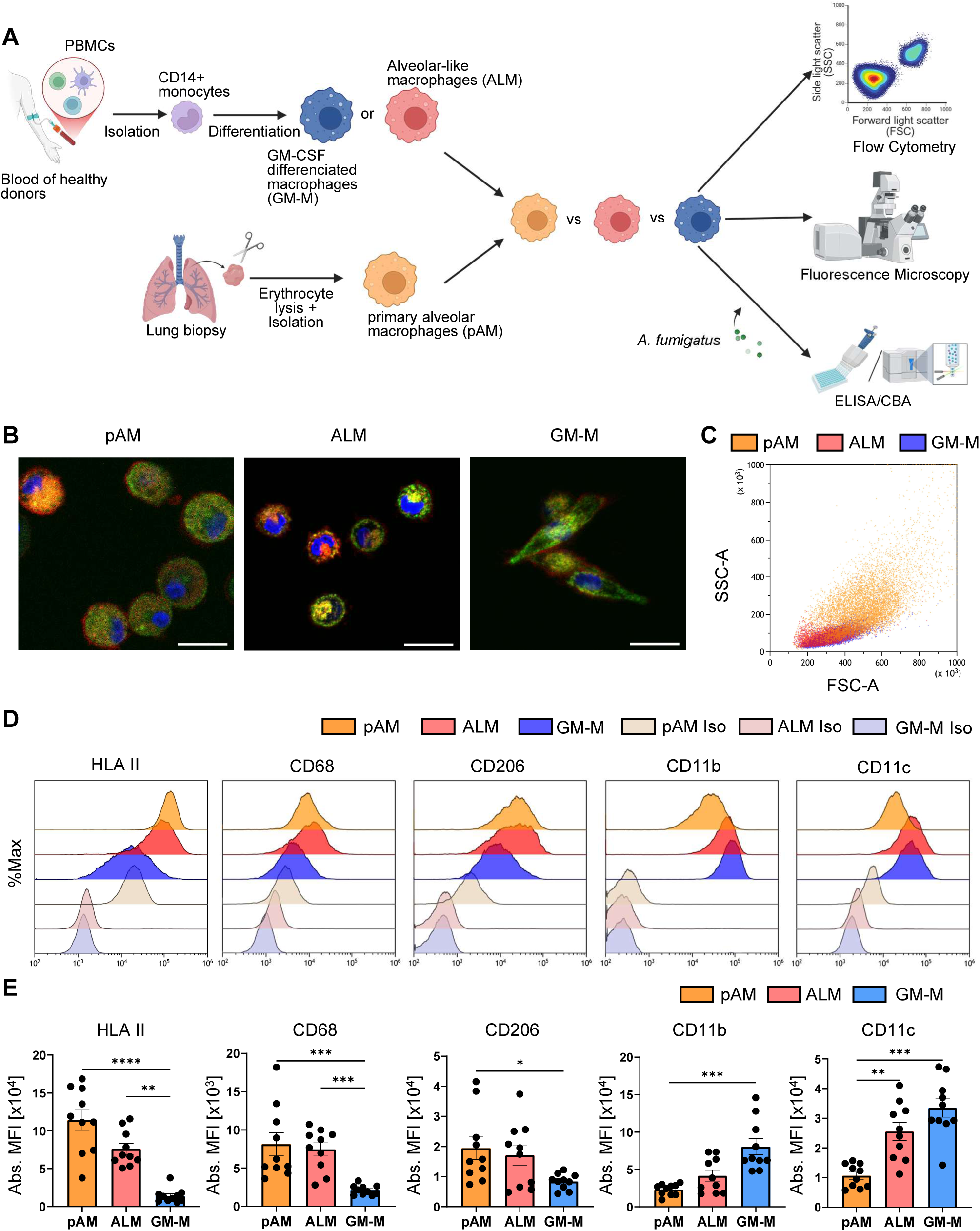
ALM resemble pAM in steady-state. (A) Schematic visualization of assays used to compare pAMs to ALMs and GM-Ms. Created with Biorender.com (B) Phenotypic evaluation of pAM, ALM and GM-M using fluorescence microscopy. The pictures were taken on a ZEISS CLSM780 with a 40x objective and cropped for better visualization. Scale bar = 20 µm (C) pAM, ALM and GM-M show differences in size and granularity in flow cytometry visualized by the SSC-A and FSC-A. (D) Flow cytometry analysis of pAM markers selected from Bain and MacDonald 2022 *Mucosal Immun.* Cells were stained with indicated antibodies or the corresponding isotype controls. Representative histograms visualize mean fluorescence intensity (MFI) of individual Mφ population markers as well as background controls. (E) Quantification of the MFI measured by flow cytometry analysis. n = 10 +/-SEM, Kruskal-Wallis-test, Dunn‘s Multiple Comparison; * p &< 0.05, ** p &< 0.01, *** p &< 0.001, **** p &< 0.0001

Having established that ALMs closely resemble pAMs under steady-state conditions, we next evaluated functional responses to *A. fumigatus* infection by measuring cytokine secretion (Fig. 2A). Following *A. fumigatus* ATCC46645 infection at MOI 1 for 6h, both pAMs and ALMs exhibited robust induction of cytokines (Fig. 2B, Fig. S2A) in contrast to GM-Ms. This was particularly evident for TNF, CCL3, CCL4, IL-1b, and IL-8. Additionally, ALMs showed significant induction of CCL5, CXCL9, CXCL10, and CXCL11 relative to uninfected controls, whereas pAMs displayed higher baseline secretion of these cytokines. GM-Ms exhibited limited responsiveness, with significant induction observed solely for CXCL9, alongside significantly elevated baseline levels of IL-1b, IL-8, and CCL5 compared to ALM. Increase of fungal burden (MOI 5) and extended incubation (9h) displayed maintenance of elevated cytokine response in pAM and ALM infected cells (Fig. 2C), particularly for TNF, CCL3, and CCL4. Comparison between pAMs and GM-Ms or ALMs and GM-Ms furthermore displayed significant differences in the TNF or CCL4 cytokine levels secreted by infected cells. In addition, GM-M failed to secrete CCL8, CCL20, IL-6, or CXCL10 while ALM exhibited increased CXCL10 secretion, which we did not observe for pAMs due to high background secretion. No substantial differences were observed for CCL2 (Fig. S2B). Collectively, our data indicates that ALMs share key phenotypic and functional characteristics with pAMs under both steady-state and *A. fumigatus* infection conditions.

**Figure 2:**
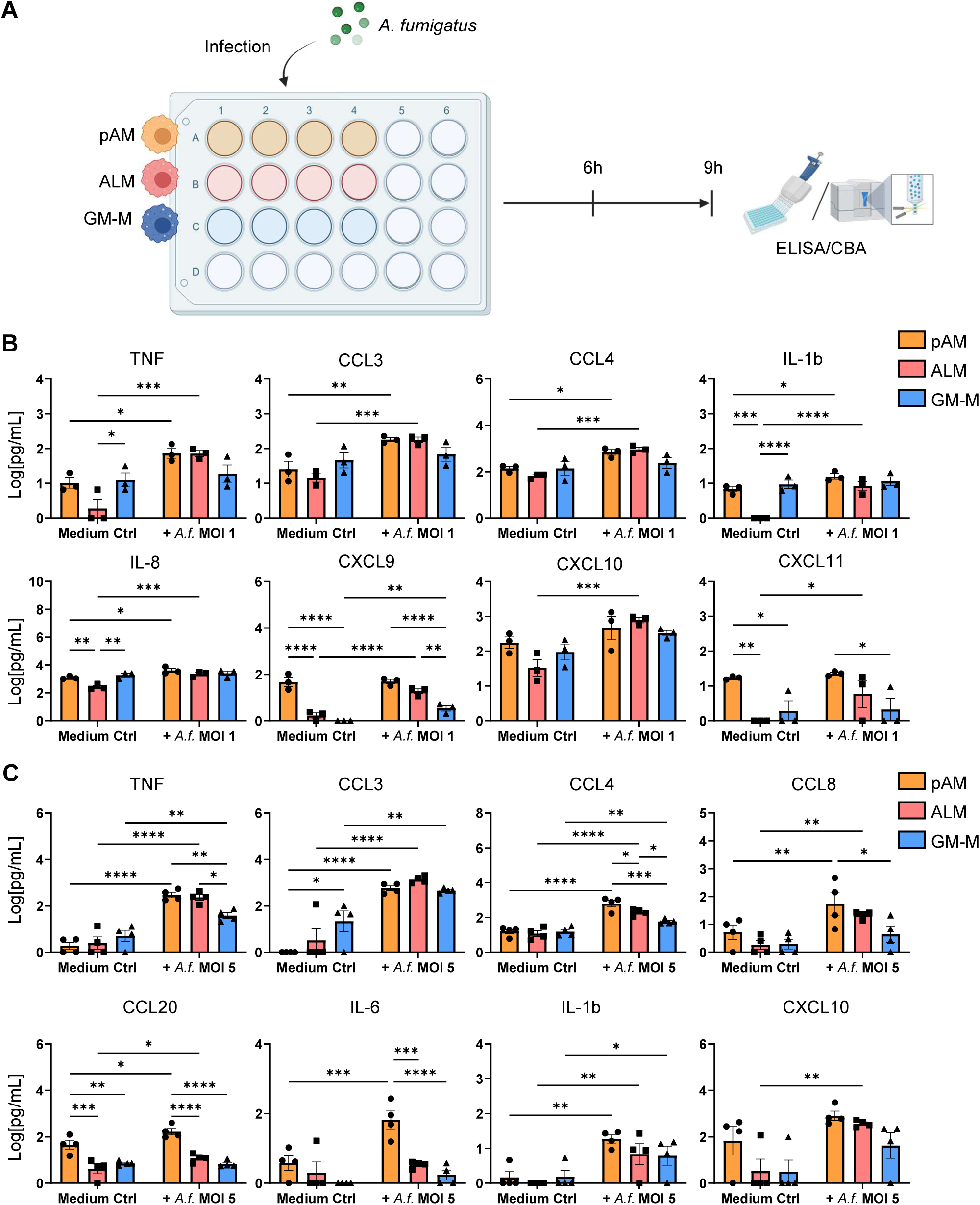
ALM resemble pAM during *Aspergillus fumigatus* infection regardless of fungal burden or infection time. (A) Schematic visualization of functional assays. pAM, ALM and GM-M were infected with *A. fumigatus* ATCC46645 at an MOI 1 for 6h or at an MOI 5 for 9h. Figure created with Biorender.com (B) Cytokine secretion of pAM, ALM and GM-M after infection with *A. fumigatus* ATCC46645 MOI 1 for 6h. Data was transformed using log(Y+1). Cytokines were measured by employing a custom ProcartaPlex assay kit. n = 3 +/-SEM, Ordinary One-Way ANOVA, Tukey‘s Multiple Comparison, * p &< 0.05, ** p &< 0.01, *** p &< 0.001, **** p &< 0.0001 (C) Evaluation of Mφ secreted cytokines upon infection with *A. fumigatus* ATCC46645 MOI 5 for 9h. The data was transformed using log(Y+1). n = 4 +/-SEM, Two-Way-ANOVA, Tukey‘s Multiple Comparison, * p &< 0.05, ** p &< 0.01, *** p &< 0.001, **** p &< 0.0001

### 3.2 ALM efficiently phagocytose *A. fumigatus* conidia

Given the observed phenotypic and functional distinctions between Mφ models, we aimed to further characterize the functional capacity of ALMs in comparison to GM-Ms. GM-Ms have been widely used to investigate host-pathogen interactions with *A. fumigatus* (Gonçalves et al. 2020; Günther et al. 2024; Brummer, Maqbool, and Stevens 2001), providing a valuable reference framework for our analyses. Building on this foundation, we evaluated key Mφ effector functions, including phagocytosis and intracellular killing, central for antifungal defense (Luther et al. 2008). We employed the *A. fumigatus* fluorescent reporter strain (FLARE), which combines a DsRed signal to indicate conidial viability with a fluorescent tracer that persists after conidial death. Following phagolysosomal uptake of conidia, the DsRed signal diminishes within approximately 45 min, thereby serving as a proxy for intracellular conidial killing (Jhingran et al. 2012; Maselli, Laevsky, and Knecht 2002). Representative gating strategies and microscopy images are provided in the supplement (Fig. S3A & B).

As outlined in Fig. 3A, we infected ALMs and GM-Ms with FLARE-labelled conidia at MOIs ranging from 0.5 to 3 and analyzed uptake and killing at 2h and 6h post-infection. The time points were selected as phagocytosis in murine Mφ was proven to last up to 2h and killing begins after 6h of phagocytosis (Philippe et al. 2003). We distinguished extracellular from internalized conidia using CFW staining. Quantitative analysis revealed that ALMs phagocytosed conidia significantly more rapidly and efficiently than GM-Ms across all tested conditions (Fig. 3B). Despite these differences in uptake kinetics, both Mφ populations exhibited comparable killing efficiencies. Within ALMs, however, killing efficiency varied depending on fungal load and increased over time. Time-course analysis further demonstrated that ALMs progressively enhanced conidial uptake, particularly at higher MOIs (Fig. 3C), indicating sustained phagocytic activity under increased fungal challenge. Consistently, intracellular killing by ALMs significantly increased over time across all conditions.

**Figure 3:**
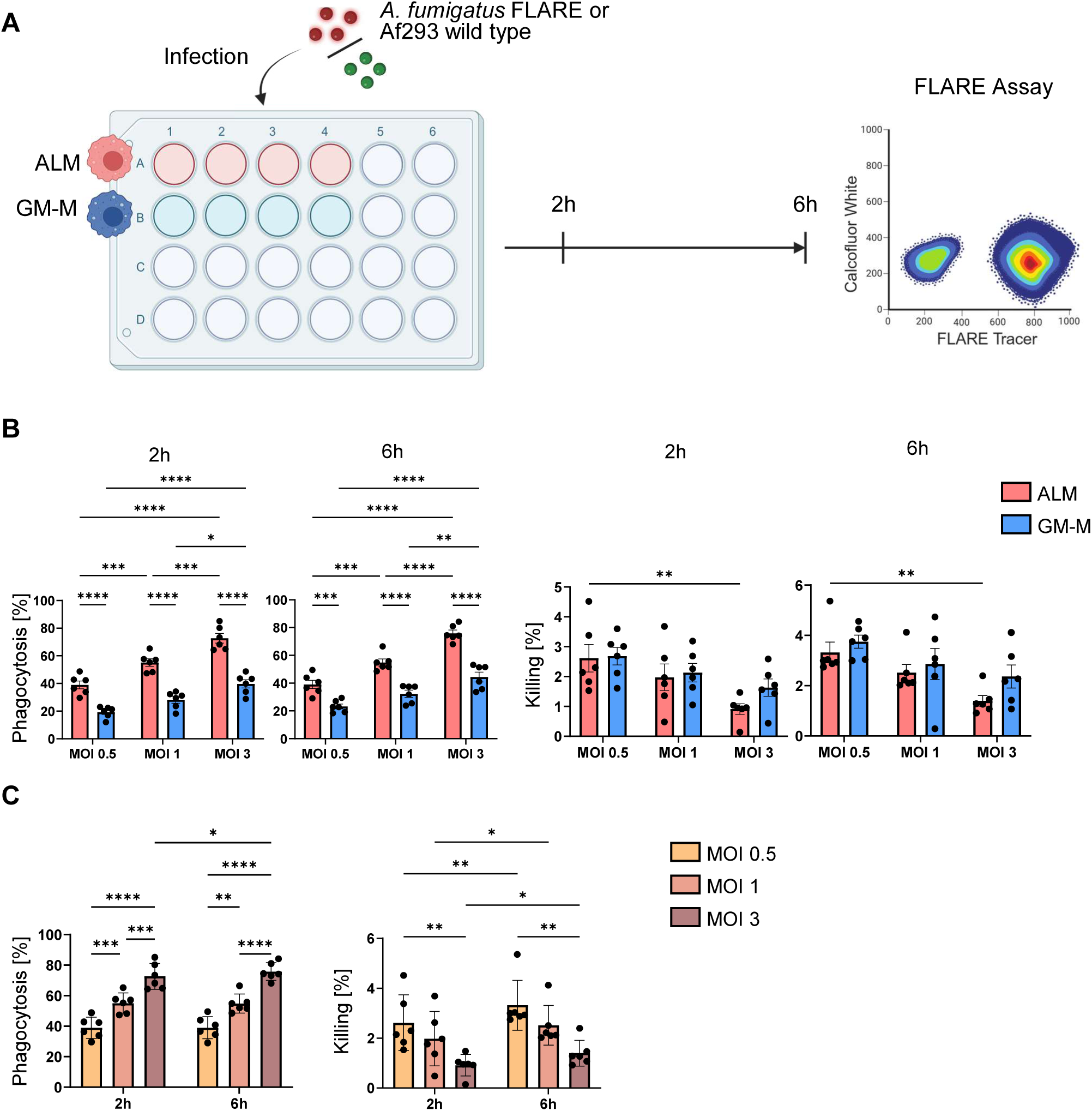
ALMs exhibit fast and efficient phagocytosis of *A. fumigatus*. (A) Visual representation of the FLARE infection setup. Mφs infected with *A. fumigatus* Af293 MOI 3 were used as isotype control to distinguish between positive staining and background signals as well as killed and live conidia. Created with Biorender.com (B) Evaluation of FLARE infected ALM and GM-M. ALM and GM-M were infected with an MOI 0.5-3 for 2h and 6h respectively before counterstaining with calcofluor white (CFW). Phagocytosis evaluated the percentage of macrophages which have phagocytosed any conidia. Killing efficacy is displayed as percentage of macrophages which contain killed conidia from all macrophages which phagocytosed conidia. n = 6 +/-SEM, Two-Way ANOVA, Tukey’s Multiple Comparison, * p &< 0.05, ** p &< 0.01, *** p &< 0.001, **** p &< 0.0001 (C) Time dependent differences in phagocytic and killing efficacy of ALM cells. n = 6 +/-SEM, Two-Way ANOVA, Tukey’s Multiple Comparison, * p &< 0.05, ** p &< 0.01, *** p &< 0.001, **** p &< 0.0001

GM-Ms also displayed time-dependent increases in conidial uptake, although at lower overall levels compared to ALMs, and showed only modest increases in killing efficiency (Fig. S3C). In summary, these findings demonstrate that ALMs combine rapid and efficient phagocytic uptake with effective intracellular killing of *A. fumigatus* conidia, consistent with key functional properties attributed to pAMs in the lung (Luther et al. 2008).

### 3.3 ALM mount a rapid and robust pro-inflammatory transcriptional response to *A. fumigatus*

Building on the observation that ALMs rapidly phagocytose *A. fumigatus* conidia, we next sought to characterize the transcriptional response to fungal challenge. To this end, we performed dual RNA-sequencing using ALMs and GM-Ms generated from matched donors. Both Mφ populations were infected with *A. fumigatus* conidia (MOI 5) for 6h and 9h, alongside uninfected controls to simultaneously assess host defense and fungal counter-defense mechanisms (Fig. 4A) related to variations in the fungal morphotype. Principal component analysis (PCA) clearly separated all experimental conditions based on infection status and Mφ differentiation (Fig. 4B). Upon infection, ALMs underwent pronounced transcriptional reprogramming at both time points, with 243 differentially expressed genes (DEGs) at 6h and 438 DEGs at 9h (Fig. S4A, Fig. 4C). Visualization of infection- and activation-associated genes (Margalit and Kavanagh 2015; van de Veerdonk et al. 2017) demonstrated that ALMs maintained a homeostatic profile in uninfected conditions, whereas infection triggered a strong induction of pro-inflammatory mediators (Fig. 4D). These include TNF, CCL3, CCL4, IL-8, CXCL9, and CXCL10. In addition, ALMs upregulated immune regulatory markers such as CD274 and CD83 and the adhesion molecule ICAM1. Pathway analysis further revealed that ALMs undergo metabolic reprogramming upon infection, characterized by suppression of oxidative phosphorylation at both 6h and 9h (Fig. S4B, Fig. 4E), alongside activation of canonical inflammatory pathways, including NF-kB- and TNF-signaling, as well as pathways associated with cell recruitment (Fig. 4E).

**Figure 4:**
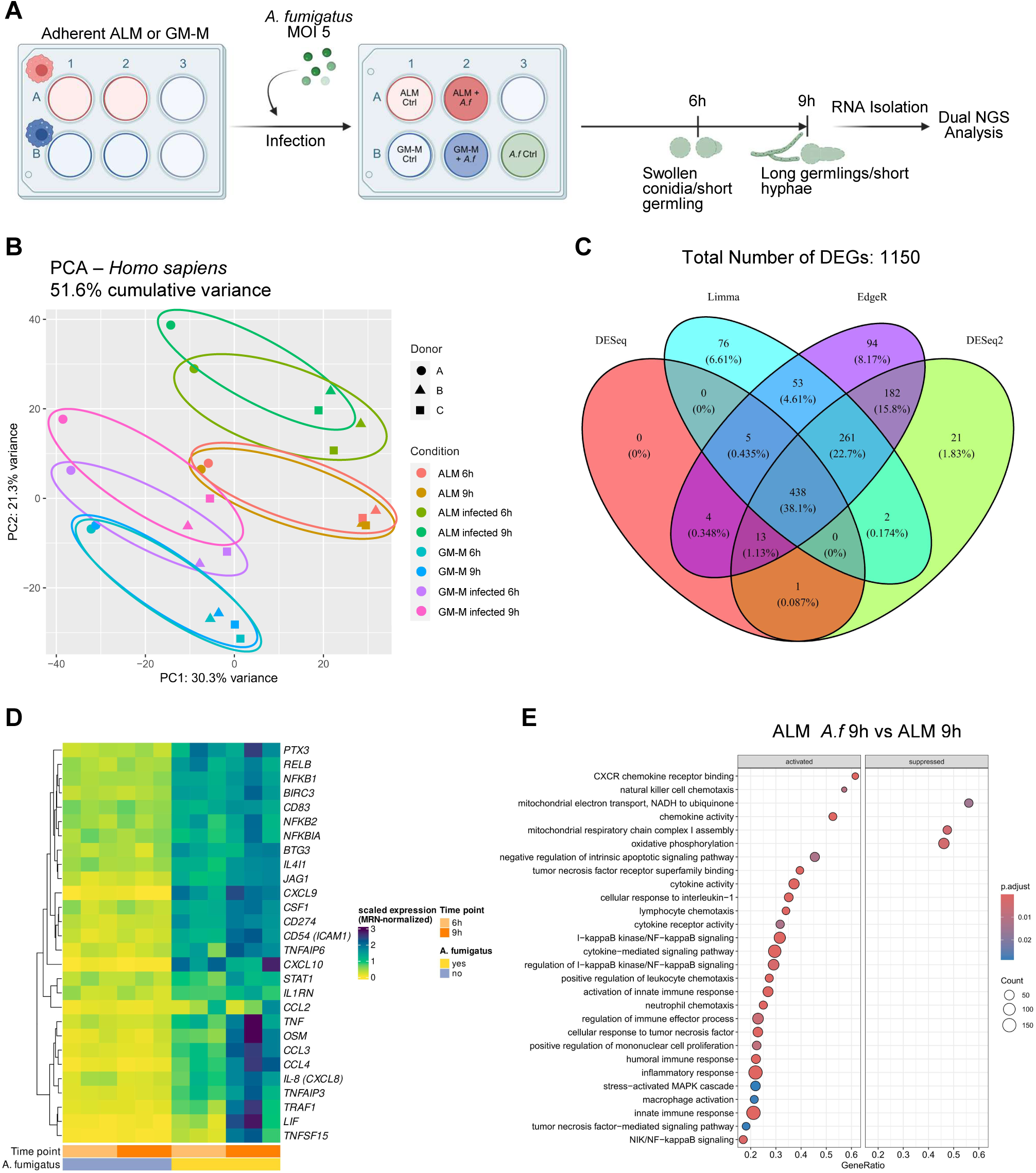
ALMs react with a fast upregulation of pro-inflammatory genes and respective pathways upon *A. fumigatus* challenge. (A) Representative figure to visualize the infection setup used for the dual RNA-sequencing analysis. Created with Biorender.com (B) PCA plot visualizing donor (n = 3) and condition variances for the human transcriptome after quality control. (C) Venn diagram of ALMs infected for 9h with *A. fumigatus* ATCC46645 MOI 5. (D) Heatmap highlighting the expression level of infection relevant genes of ALM uninfected or infected for 6h and 9h with *A. fumigatus* ATCC46645 MOI 5. (E) Pathway analysis of ALMs infected for 9h with *A. fumigatus* ATCC46645 MOI 5. ALM cells quickly downregulate genes for oxidative phosphorylation and further upregulate pro-inflammatory pathways like NF-kB- or TNF-signaling at 9h of infection.

GM-Ms also responded to infection but exhibited delayed and gradual transcriptional adaptation. At 6h post-infection, GM-Ms showed a limited response with only 16 DEGs, which increased to 283 DEGs at 9h (Fig. S4C). Elevated baseline expression of inflammatory genes in uninfected GM-Ms indicates a pre-activated state, which may influence their response dynamics (Fig. S4D). By 9h, GM-Ms upregulated cytokine- and chemokine-associated pathways. However, activation of NF-kB- and TNF-signaling, as well as immune cell recruitment-related pathways, remained less pronounced compared to ALMs (Fig. S4E). Like ALMs, GM-Ms exhibited suppression of oxidative phosphorylation.

We validated these transcriptional findings at the protein level using multiplex cytokine analysis of the three initial donors (Fig. 5A, Fig. S5). ALMs significantly secreted TNF, CCL3, CCL4, IL-1b, and IL-6 as early as 6h, with increase at 9h. In contrast, GM-M displayed delayed cytokine production and lower overall secretion levels, even at 9h post-infection. Validation in an expanded cohort of six donors (Fig. 5B) confirmed these patterns, with ALMs consistently producing higher levels of TNF, CCL3, CCL4, and CXCL10 than GM-M, particularly at 9h of fungal challenge. Collectively, these results demonstrate that ALMs mount a rapid, coordinated and robust pro-inflammatory response to *A. fumigatus,* highlighting their value as a physiologically relevant *in vitro* model for studying early host-pathogen interactions.

**Figure 5:**
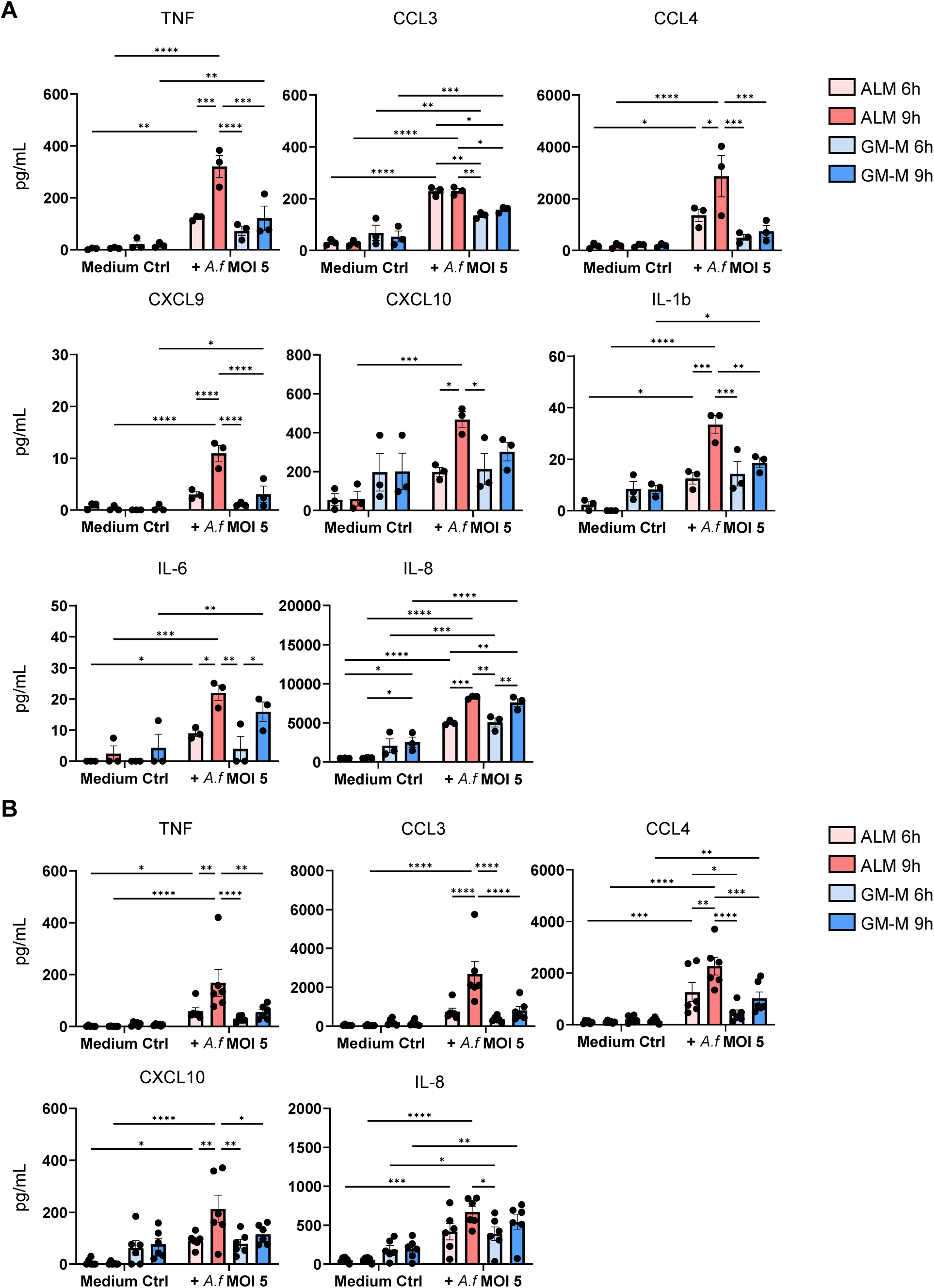
ALMs secrete significantly more cytokines than the typical blood-derived GM-Ms. (A) Quantification of cytokine secretion in supernatant of ALM and GM-M donors used for dual RNA-sequencing analysis. ALM and GM-M were left uninfected or infected with *A. fumigatus* ATCC46645 MOI 5 for 6h and 9h. The cytokines were measured employing a custom ProcartaPlex assay kit. n = 3 +/-SEM, Two-Way ANOVA, Tukey‘s Multiple Comparison;* p &< 0.05, ** p &< 0.01, *** p &< 0.001, **** p &< 0.0001 (B) Validation of the dual RNA-sequencing dataset by singleplex ELISA including an expanded cohort. n = 6 +/-SEM, Two-Way ANOVA, Tukey‘s multiple comparison; * p &< 0.05, ** p &< 0.01, *** p &< 0.001, **** p &< 0.0001

### 3.4 *A. fumigatus* transcriptome analysis mounts an enhanced counter-defense response during interaction with ALMs

To complement our analysis of host responses, we next investigated fungal transcriptional adaptations during Mφ interaction (Fig. 6A). PCA revealed clear clustering of fungal samples according to Mφ type and infection time point, indicating distinct transcriptional programs under each condition (Fig. 6B). During interaction with ALMs, *A. fumigatus* underwent extensive transcriptional reprogramming, particularly at 9h, with 2,325 DEGs (Fig. 6C). Compared to unchallenged controls, the fungus strongly upregulated genes associated to stress adaptation and immune evasion, including *cat1* (Afu3g02270) (Abad et al. 2010; Keizer et al. 2022) or key components of the gliotoxin biosynthetic pathway (*gliZ*/Afu6g09630, *gliA*/Afu6g09710, *gliG*/Afu6g09690) (Knowles et al. 2020). Additionally, *A. fumigatus* induced allergen-encoding genes such as *aspf1* (Afu5g02330) and *aspf2* (Afu4g09580), with *aspf1* showing significant induction during interaction with ALMs (Fig. 6D). Pathway analysis revealed a dynamic transcriptional response characterized by suppression of growth-associated processes, including ergosterol biosynthesis and nuclear division (Alcazar-Fuoli et al. 2008), alongside activation of transcriptional regulation pathways at 6h (Fig. 6E, left panel). At 9h, the fungal response shifted toward activation of virulence-associated programs, including gliotoxin production, secondary metabolite biosynthesis, and IgE binding pathways (Fig. 6E, right panel), indicating an intensified adaptive fungal response to ALM-mediated immune pressure.

**Figure 6:**
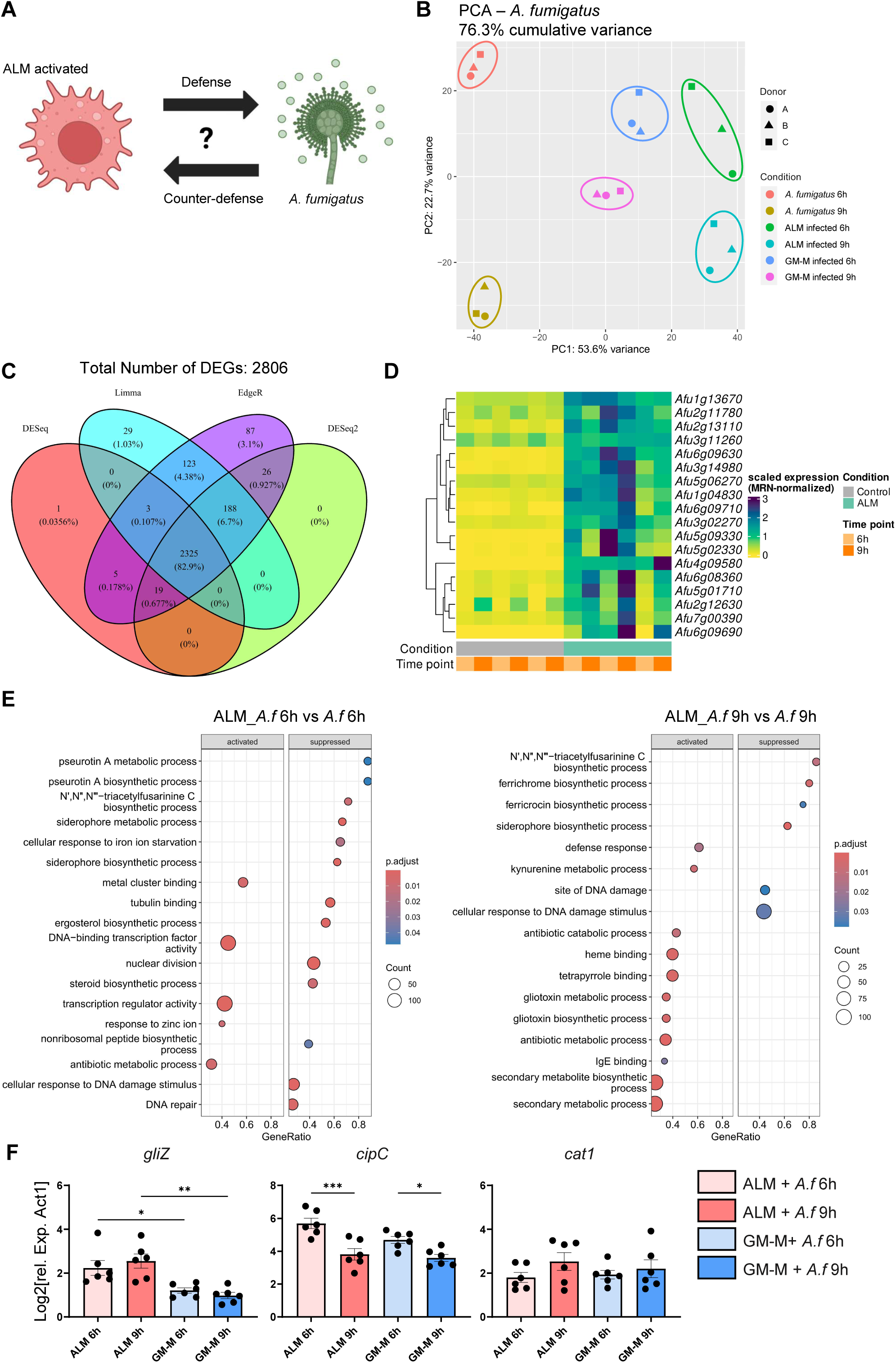
Analysis of the fungal transcriptome reveals a strong counter-defense response of *A. fumigatus* upon ALM stimulation used for immune evasion. (A) Graphical illustration to visualize the idea behind the dual RNA-sequencing analysis to compare both human and fungal transcriptomic profiles. Created with Biorender.com (B) *A. fumigatus* PCA plot of the dual RNA-sequencing analysis visualizing differences in the transcriptomic profile. (C) Venn diagram of fungal DEGs after 9h of challenge with ALM. (D) Heatmap of respective virulence genes expressed by *A. fumigatus* in contact to ALM next to the respective unchallenged control. (E) Representative pathways regulated by *A. fumigatus* transcriptome after 6h and 9h of ALM encounter. The dot plot visualizes pathways activated or suppressed by *A. fumigatus* ATCC46645 upon ALM challenge. (F) Validation of selected genes upregulated in *A. fumigatus* upon macrophage challenge. n = 6 +/-SEM, Ordinary One-Way ANOVA, Šidák‘s Multiple Comparison; * p &< 0.05, ** p &< 0.01, *** p &< 0.001

During interaction with GM-Ms, *A. fumigatus* also adjusted its transcriptional program but to a lesser extent. At 9h, fewer DEGs were detected (1,331 DEGs; Fig. S6B), and induction of virulence-associated genes was comparatively reduced (Fig. S6A). While suppression of growth-related pathways was also evident at 6h (Fig. S6C), GM-M interaction preferentially induced pathways associated with biofilm formation, which were not prominently activated during ALM interaction. Notably, key pathways such as gliotoxin biosynthesis and kynurenine metabolism were not significantly induced in response to GM-Ms. We validated these dual RNA-sequencing results by qRT-PCR analysis of selected fungal genes (*gliZ*, *cat1*, and *cipC*), confirming the observed transcriptomic profiles at both time points (Fig. 6F).

Together, these results indicate that *A. fumigatus* mounts a more pronounced and coordinated counter-defense response during interaction with ALMs. This enhanced host-pathogen interplay not only reflects the stronger immune activation elicited by ALMs but also facilitates the identification of fungal adaptation mechanisms, including the induction of genes such as *aspf1*, that may be less apparent in alternative Mφ models such as GM-Ms.

### 3.5 Host-pathogen interaction networks highlight coordinated ALM and *A. fumigatus* infection responses

To further characterize the bidirectional interaction between ALMs and *A. fumigatus*, we constructed predicted host-pathogen protein-protein interaction (PPI) networks based on the dual transcriptomic dataset as described in the methods section. Therefore, we prioritized candidate proteins based on features associated with host-pathogen interactions, including predicted GPI-anchors, transmembrane domains, signaling peptides, secretion signals and subcellular localization. PPI networks were computed using STRING DB. Consistent with the pathway enrichment analyses above, network analysis revealed a persistent modulation of *A. fumigatus* interaction on the host aerobic electron transport chain-associated proteins at both investigated time points (Figure 7A & B). In parallel, ALMs robustly modulated a central inflammatory network enriched for TNF signaling-associated proteins, including ICAM1, TLR2, CSF1, LILRB1, and CXCL10. Employing the STRING DB enrichment module, we found that ALMs exhibited a cluster visualizing a pronounced CXCL10-associated inflammatory signature at 6h post-infection, whereas expression of both CSF2 receptor subunits became more prominent at 9h, indicating sustained immune activation and macrophage responsiveness. On the fungal side, the predicted interaction landscape initially emphasized proteins associated with galactose metabolism and protein translation at 6h post-infection. By 9h, however, the fungal network shifted toward a pronounced counter-defense response signature characterized by proteins linked to stress adaptation, respirasome-associated proteins, and responses to reactive oxygen species, suggesting adaptation to increasing host-imposed immune pressure. Notably, the analysis revealed several established virulence-associated proteins, including the antioxidant enzymes catalase Cat1 and superoxide dismutase SodB. Collectively, these host-pathogen interaction networks further support the concept that ALMs mount a coordinated inflammatory response that induces extensive fungal counteradaptation during early infection.

**Figure 7:**
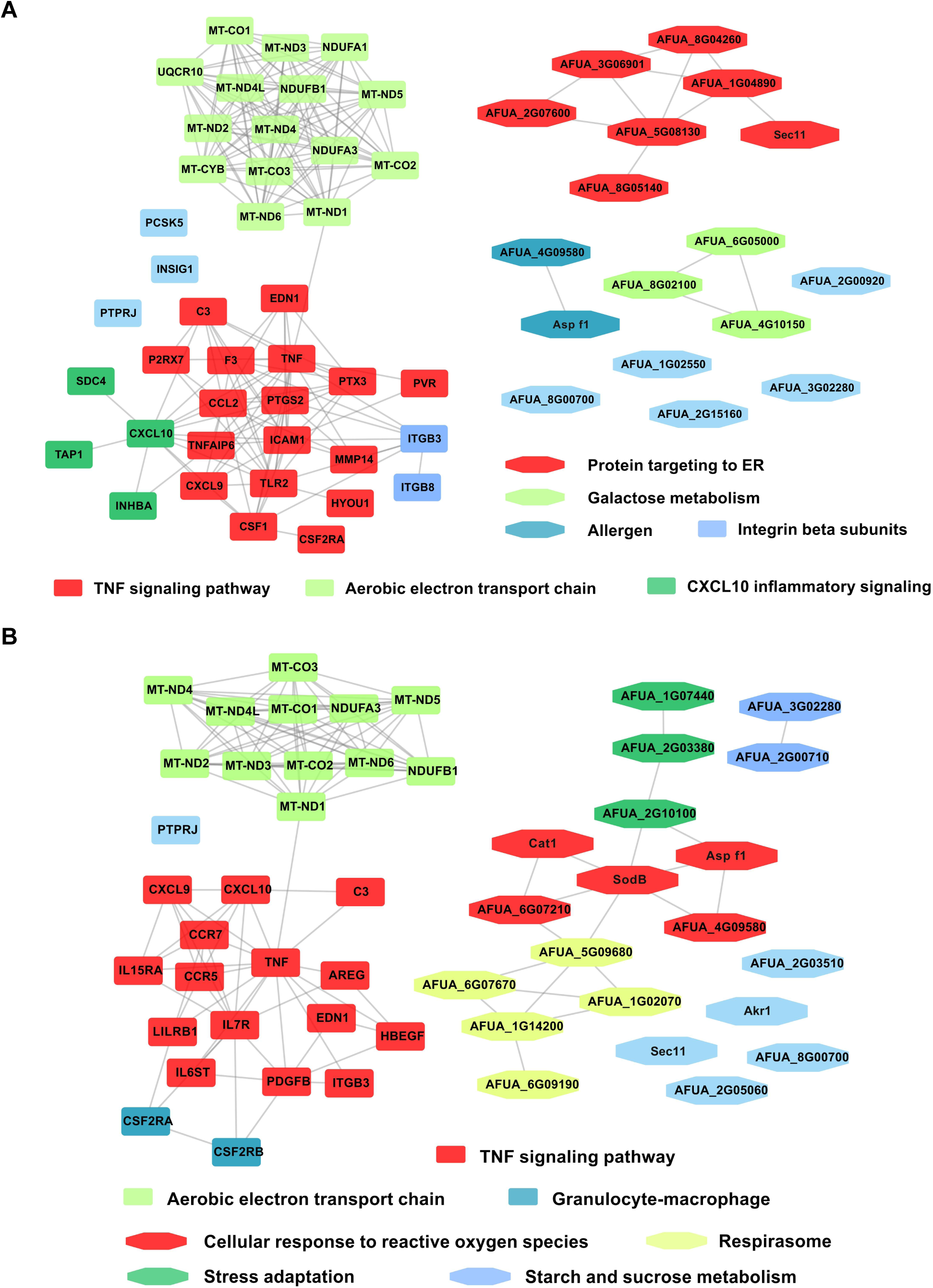
Host–pathogen protein–protein interaction (PPI) networks generated from the dual RNA-sequencing dataset. The plots display the PPIs (A) 6h and (B) 9h. The networks were constructed as indicated in the methods section. Modules and interactions were identified using STRING DB. Node shapes distinguish host proteins from pathogen-associated proteins, whereas edges indicate interactions between or functional influence on the proteins.

### 3.6 Fungal allergens contribute to immune evasion during macrophage interaction

The induction of allergen-encoding genes *aspf1* and *aspf2* in our dual RNA-sequencing analysis prompted us to investigate their functional role during early Mφ interactions. We infected pAMs, ALMs, and GM-Ms with *A. fumigatus* wild type (WT; strain CEA17 Δ*akuB KU80*) and the corresponding knockout mutant strains Δ*aspf1* and *Δaspf2* for 9h. Subsequently, we quantified cytokine and chemokine secretion using cytometric bead array (CBA) analysis.

Consistent with our previous observations, pAMs and ALMs mounted robust cytokine responses upon infection, whereas GM-Ms showed limited responsiveness relative to uninfected controls (Fig. 8A). Elevated baseline cytokine levels in GM-M, including CCL2, CCL3, CXCL10, and IL-8 further support their pre-activated phenotype. Comparative analysis of WT and mutant strains revealed largely comparable cytokine responses across all three Mφ populations. However, infection with the Asp f1-deletion mutant resulted in enhanced cytokine induction relative to uninfected controls for several mediator including IL-1b, IL-10, CCL4, CCL8, CXCL10, TNF, IL-6, or IL-8, depending on the Mφ type (Fig. 8A & B, Fig. S7A). Notably, GM-Ms displayed significant induction of TNF and CXCL10 only in response to the Asp f1-deficient strain, but not to WT infection, suggesting that Asp f1 may modulate host inflammatory responses.

**Figure 8:**
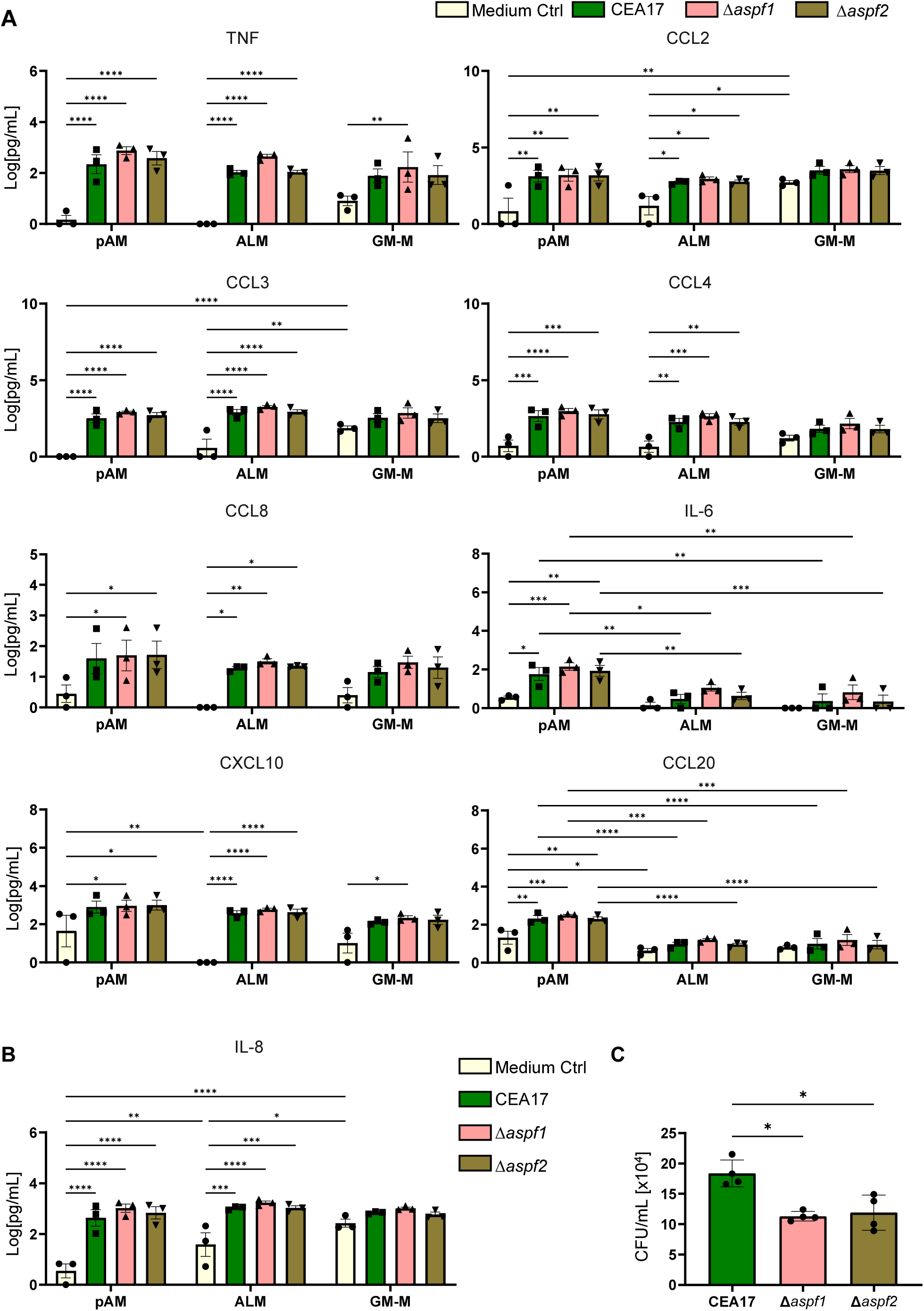
pAM and ALM show strong infection responses upon challenge with *A. fumigatus* CEA17, Δ*aspf1* and Δ*aspf2*. (A) Cytokine secretion analysis of pAM, ALM and GM-M infected with MOI 5 of *A. fumigatus* CEA17 or its respective knockout mutants after 9h. Cytokines were analyzed using a custom LEGENDplex CBA kit. The values were transformed using log(Y+1). n = 3 +/-SEM; Two-Way ANOVA, Tukey’s Multiple Comparison; * p &< 0.05, ** p &< 0.01, *** p &< 0.001, **** p &< 0.0001 (B) Singleplex ELISA analysis of IL-8 secreted by pAM, ALM or GM-M upon *A. fumigatus* CEA17, Δ*aspf1* and Δ*aspf2* MOI 5 infection. n = 3 +/-SEM; Two-Way ANOVA, Tukey’s Multiple Comparison; * p &< 0.05, ** p &< 0.01, *** p &< 0.001, **** p &< 0.0001 (C) Colony forming unit analysis of ALMs infected with MOI 5 of *A. fumigatus* CEA17 or its respective knockout mutants after 9h. ALMs were lysed with cold ddH_2_O and the conidia suspension was plated on malt agar plates at 37°C for 18h. n = 4 +/-SEM, RM-ANOVA, Dunnett’s Multiple comparison; * p &< 0.05

To further assess the functional impact of allergen deficiency, we quantified fungal survival following infection of ALMs using colony-forming unit (CFU) assays. Both Δ*aspf1* and *Δaspf2* showed significantly reduced CFU recovery compared to WT (Fig. 8C), indicating enhanced Mφ-mediated fungal killing by ALMs in the absence of these allergens. To exclude confounding effects on fungal germination or uptake, control experiments confirmed that these differences were not attributable to altered fungal growth or phagocytic uptake (Fig. S7B-D). Together, these findings indicate that the allergens Asp f1 and Asp f2 contribute to fungal persistence during Mφ interaction, at least in part by contributing to immune evasion. Their absence enhances Mφ-mediated killing in ALMs, and modestly augments cytokine responses, emphasizing their role in shaping host-pathogen dynamics.

## 4 Discussion

Invasive fungal infections including IPA remain a major cause of morbidity and mortality in immunocompromised patients, particularly when diagnosis is delayed or insufficiently sensitive (Elalouf, Elalouf, and Rosenfeld 2023; Kousha, Tadi, and Soubani 2011; Wichmann et al. 2025). The environmental mold *A. fumigatus* deploys multiple highly specialized immune evasion strategies, including masking of pathogen-associated molecular patterns by melanin and the hydrophobin RodA, resistance to oxidative stress, and rapid germination at physiological temperatures of the host (Liu et al. 2024; Heinekamp et al. 2015). Upon inhalation, pAMs serve as the first line of defense by encountering inhaled conidia, thereby maintaining pulmonary homeostasis and initiating antifungal immunity (Balloy and Chignard 2009; Yan et al. 2025). However, the early cellular and molecular determinants impacting fungal clearance versus progression to invasive disease remain insufficiently understood, in part due to the lack of physiologically relevant *in vitro* models.

In the present study, we adapted and systematically validated a previously established ALM model derived from human primary monocytes and demonstrate that these cells closely recapitulate key phenotypic and functional features of pAMs. Consistent with previous work (Pahari et al. 2023), ALMs exhibited a round morphology and expressed canonical Mφ markers, closely resembling pAMs, whereas GM-Ms displayed a divergent phenotype. Notably, the expression of MHC II molecules and the scavenger receptor CD68, key mediators of antigen presentation and phagocytic activity (Chen and Jensen 2008; Chistiakov et al. 2017; Zhang et al. 2022), aligned closely between ALMs and pAMs, while GM-Ms differed across all assessed markers. Functionally, ALMs mirrored pAM-like cytokine responses upon *A. fumigatus* challenge, independent of fungal load and infection time. Specifically, TNF, CCL3, IL-1b, and IL-8, hallmark mediators of antifungal immunity (Shankar et al. 2024) were significantly induced in ALM and pAM, whereas GM-M exhibited elevated baseline cytokine levels, indicative of a pre-activated state. These findings demonstrate that ALMs maintain a homeostatic steady-state under resting conditions while mounting a rapid inducible response upon pathogen challenge, characteristic of tissue-resident Mφs (Mosser, Hamidzadeh, and Goncalves 2021; Sun, Cui, and Zhang 2026). In contrast, GM-Ms appear constitutively primed, likely due to GM-CSF-driven differentiation (Lotfi et al. 2020; Petrina et al. 2024). Together, these results highlight how differentiation context shapes Mφ identity and emphasize the limitations of the employed conventional GM-CSF-based model.

Using the FLARE-based assay, we directly quantified phagocytosis and killing of *A. fumigatus,* two critical early defense mechanisms against inhaled fungal conidia (Luther et al. 2008; Jhingran et al. 2012). ALMs internalized conidia more rapidly and efficiently than GM-Ms across all tested conditions. This behavior reflects the physiological role of alveolar Mφs, which continuously survey the airway environment and rapidly clear particulate matter (Luther et al. 2008; Neupane et al. 2020). Although, intracellular killing efficacy rates were broadly comparable, the accelerated uptake kinetics of ALMs likely provide a decisive advantage by restricting fungal germination, a key step in disease progression (Savers et al. 2016). Notably, ALMs sustained phagocytic activity under high fungal loads and increased killing over time, underscoring their functional adaptability.

At the transcriptional level, paired donor analyses revealed that ALMs undergo rapid and coordinated activation upon fungal challenge. Infection triggered metabolic reprogramming was characterized by downregulation of oxidative phosphorylation, the predominant energy-generating pathway in tissue-resident alveolar Mφs (Woods and Mutlu 2025) and consistent with established Mφ activation programs during infection (Wculek et al. 2023; Hatinguais et al. 2021; Gonçalves et al. 2020). Concurrently, ALMs strongly induced pro-inflammatory transcriptional programs, including TNF- and NF-kB-signaling pathways, alongside chemokines such as CCL3 and CXCL10 that drive immune cell recruitment (Shankar et al. 2024). These results are consistent with the established role of inflammatory signaling in antifungal host defense (Mezger et al. 2008; Guo et al. 2020). In contrast, GM-Ms responded more slowly and less synchronously, reinforcing the concept that Mφ differentiation state dictates both the timing and magnitude of antifungal immunity.

In our previous study (Zoran et al. 2019), we identified etanercept treatment and reduced monocyte counts as risk factors for IPA in a clinical setting and demonstrated that etanercept can impair Mφ-associated immune responses to *A. fumigatus*. Specifically, modulation of the TNF- and CXCL10-associated responses, as well as effects on NF-kB-related signaling pathways, suggested that disruption of key inflammatory circuits may compromise antifungal immunity. Based on these results, the present study provides mechanistic support that ALMs rapidly activate these critical pathways upon infection. In contrast, the delayed response observed in GM-Ms suggests reduced responsiveness to pathogen-specific cues. Mechanistically, the rapid and coordinated activation in ALMs likely reflects enhanced pathogen sensing and efficient signal transduction, enabling effective early immune control despite fungal immune evasion strategies.

While our past work (Zoran et al. 2019) focused primarily on human signaling responses, by integrating dual RNA-sequencing in our current study, we captured the dynamic interplay between host defense and fungal adaptation. Particularly, *A. fumigatus* mounted a strikingly stronger transcriptional counter-response when challenged with ALMs, including the induction of cytotoxic secondary metabolites like gliotoxin (Sun, Cui, and Zhang 2026) or oxidative stress defenses such as *cat1* (Abad et al. 2010; Keizer et al. 2022). This implies a tightly coupled interaction in which enhanced host immune activation directly triggers fungal adaptation mechanisms. Such bidirectional regulation was not accessible in our previous study and highlights the added value of the ALM model combined with dual transcriptomics. Markedly, the substantially stronger transcriptional response of *A. fumigatus* challenged with ALMs compared to GM-Ms included upregulation of oxidative stress defense genes such as *cat1* (Keizer et al. 2022; Abad et al. 2010), as well as genes involved in gliotoxin biosynthesis. Gliotoxin is a major virulence factor known to impair host immune function by inducing apoptosis, inhibiting phagocytosis, and suppressing NF-kB-signaling (Günther et al. 2024; Knowles et al. 2020). Furthermore, we found an upregulation of *A. fumigatus* kynurenine metabolism, a potential immune evasion mechanism to dampen the host inflammatory response (Gargaro et al. 2016; Zelante et al. 2021). The enhanced induction of gliotoxin and kynurenine biosynthesis pathways in response to ALMs indicates that the fungus may perceive these cells as a more potent immune threat, thereby activating more robust counter-defense strategies.

Analysis of the predicted PPI network further emphasizes the profound impact of *A. fumigatus* on ALM responses. The identified interaction clusters reinforced our pathway enrichment analyses and provided additional evidence that fungal challenge profoundly affects host oxidative phosphorylation, a process increasingly recognized as a central component of Mφ adaptation during fungal infection (Hatinguais et al. 2021; Gonçalves et al. 2020). In parallel, the host network prominently featured TNF-associated signaling modules, including ICAM1, TLR2, CSF1 and CXCL10, further supporting the central role of inflammatory signaling during antifungal defense and aligning with our previous work (Zoran et al. 2019). Experimental studies previously demonstrated that TNF pre-treatment enhances resistance to *A. fumigatus* in infection in neutropenic mice by increasing Mφ phagocytic activity, whereas TNF-neutralization results in elevated fungal burden and mortality in IPA mouse models (Balloy and Chignard 2009). Consistent with these observations, TLR2-mediated recognition of both conidia and hyphae induces TNF expression, while loss of TLR2 markedly impairs this response (Mambula et al. 2002; Balloy and Chignard 2009). Moreover, our previous work linked TNF-associated signaling to CXCL10 induction during *A. fumigatus* infection (Zoran et al. 2019), a relationship that was recapitulated in the current PPI network analysis. Together, these findings further support an important role of the TNF-CXCL10 axis in orchestrating early fungal immunity.

On the fungal side, the predicted interaction networks identified a distinct cluster of oxidative stress response factors, including the antioxidant enzymes Cat1 and SodB, closely associated with proteins involved in the respirasome and energy metabolism (Wu et al. 2020). During host-pathogen interaction, reactive oxygen and nitrogen species generated by Mφs impair fungal respiration and mitochondrial function (Joseph-Horne et al., 2001), necessitating efficient detoxification mechanisms. In this context, SodB expression appears to fulfill a central protective role and has additionally been implicated in energy metabolism during early fungal germination in *A. nidulans*. Consistent with these functions, deletion of *sodB* increases susceptibility to Mφ-mediated killing (Pákozdi et al., 2024; Kanamaru et al., 2025). The coordinated induction of oxidative stress defense pathways together with respiratory adaptation mechanisms therefore suggests that *A. fumigatus* actively remodels its metabolism to withstand ALM-imposed immune pressure.

Beyond classical virulence factors stated above, we identified a significant induction of allergen-encoding genes, including *aspf1* and *aspf2* as functionally relevant mediators of host-pathogen interaction upon ALM challenge. Deletion of these genes reduced fungal survival and increased susceptibility to ALM-mediated killing. Our results suggest that fungal allergens may contribute to immune evasion beyond their established role in allergic sensitization. One potential explanation is that these proteins modulate host immune signaling or interfere with intracellular killing mechanisms, thereby promoting fungal persistence. Mitogillin, encoded by *aspf1*, is a ribotoxin contributing to *A. fumigatus* pathogenicity (Liu et al. 2018) that cleaves a phosphodiester bond in the 28S ribosomal RNA, thereby inhibiting protein synthesis and inducing apoptosis (Morton et al. 2011; Lacadena et al. 2007). On the other hand, Asp f2 was identified as a central immune evasion and complement regulating protein with the ability to bind to Factor H, factor-H-like proteins and plasminogen. Hence, blocking the host innate immune response and disrupting lung epithelial cell layers to assist in early fungal infection and tissue penetration (Dasari et al. 2018). Suitable to the described role of Asp f1 as a protein biosynthesis inhibitor (Lacadena et al. 2007), our observation that Δ*aspf1* conidia induced enhanced cytokine response in our setup extend these functions by demonstrating these allergens actively modulate Mφ responses during early infection. Importantly, these fungal adaptations were substantially attenuated during interaction with GM-Ms, where fewer virulence-associated pathways were induced. This finding underscores that the choice of Mφ model not only shapes host responses but also fundamentally influences pathogen behavior, potentially masking critical virulence mechanisms in less representative systems.

Despite these advances, several limitations of our study should be considered. First, the ALM phenotype depends on defined differentiation conditions and may exhibit temporal variability, particularly after withdrawal of key factors (Pahari et al. 2023). Second, our experimental system relies on monocultures and therefore does not capture the full complexity of the alveolar niche, including epithelial interactions and multicellular immune networks. Future studies incorporating co-culture systems or organoid models will be essential to validate and extend these findings. Lastly, our computational approach did not identify direct host-pathogen protein interactions. Future studies could improve these analyses by integrating additional bioinformatic resources and interaction databases, including ortholog- and homology-based datasets derived from other fungal species like *Saccharomyces cerevisiae* or *Candida albicans* (Remmele et al. 2015). Incorporation of these resources may help to refine predicted interaction networks and advance the mechanistic understanding of alveolar Mφ-*A. fumigatus* interactions during early infection.

Collectively, our study demonstrates that ALMs provide a robust and physiologically relevant, experimentally accessible *in vitro* platform for investigating early interactions in pulmonary aspergillosis (Fig. 9). Compared to GM-Ms, ALMs more accurately reproduce the steady-state phenotype, rapid immune activation kinetics, and functional responses of pAMs. By enabling simultaneous analysis of host and pathogen programs, this model provides new insights into the dynamic balance between immune defense and fungal counter-defense. Beyond aspergillosis, ALMs offer a versatile platform for studying a broad range of respiratory infections and for identifying therapeutic targets that enhance host immunity.

**Figure 9:**
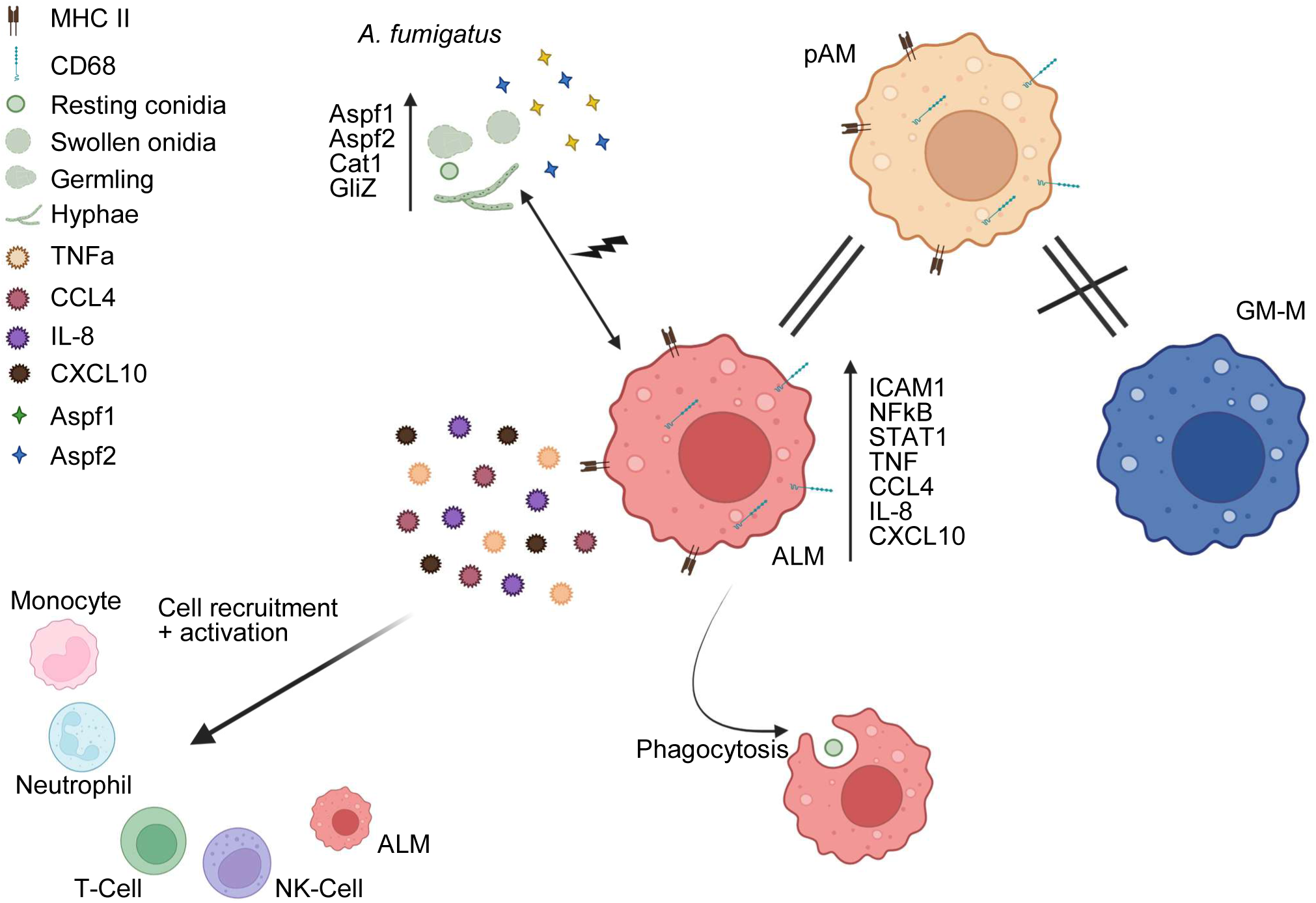
ALM are an efficient pAM resembling *in vitro* model suitable to study *A. fumigatus* infection. Summary displaying the results found in the research study. ALM resemble a pAM phenotype in steady-state as well as during infection, which we did not find for GM-M. Upon *A. fumigatus* infection, ALMs rapidly phagocytose *A. fumigatus* conidia as well as upregulate pro-inflammatory genes including cytokines and chemokines, to attract other immune cell types like neutrophils and monocytes to eliminate the fungus. On the fungal side, *A. fumigatus* encounters ALM as a important immune threat and upregulates genes encoding immune evasion mechanisms including gliotoxin and the allergens Asp f1 and Asp f2 for fungal survival. Created with Biorender.com

## 6 Author Contributions

The study was conceptualized by J.S., A.B., and J.L. Experiments were planned and performed by J.S., K.H., Z.A., D.S., M.S., A.B., and J.L. Data was analyzed by J.S., Sa.S., J.P.S., and T.D. Data were visualized by J.S., Sa.S. and J.P.S. Project administration and supervision were led by D.S., M.S., H.E., T.D., A.B., and J.L. Funding was acquired by A.B. and J.L. The original draft was written by J.S., Sa.S., J.P.S., and J.L. All co-authors reviewed, edited, and approved the manuscript.

Further information and requests for resources and reagent should be directed to and will be fulfilled by the lead contact, Prof. Dr. Juergen Loeffler.

## 8 Acknowledgments

We thank Prof. Dr. Tobias M. Hohl (Infectious Disease Service, Department of Medicine, Memorial Sloan Kettering Cancer Center, New York, NY, USA) for providing the *A. fumigatus* FLARE conidia. Furthermore, we thank Dr. Thorsten Heinekamp and Prof. Dr. Axel A. Brakhage (Molecular and Applied Microbiology, Leibniz Institute for Natural Product Research and Infection Biology, Hans Knöll Institute (HKI), Jena, Germany) for providing *A. fumigatus* CEA17 Δ*akuB KU80*, Δ*aspf1* and Δ*aspf2* conidia. We are grateful for Prof. Dr. Frank Ebel (Department of Bacteriology and Mycology, Ludwig-Maximilian University Munich, Munich, Germany) for providing *A. fumigatus* ATCC46645 conidia. Lastly, we thank the Core Unit SysMed at the University of Wuerzburg for excellent technical support, dual RNA-seq data generation and processing. We are grateful for Prof. Dr. Tom Gräfenhan (CU SysMed) for the constructive discussion and NGS expertise. This work was supported by the IZKF at the University of Wuerzburg (Project Z-6). The authors thank all blood and lung tissue donors.

Figures Fig. 1A, 2A, 3A, 4A, 6A and 9 were created in Biorender.com and can be found under the designated agreement numbers (*LZ29NG8OVH, NT29NG9PY5, RT29NGA2CL, OO29NGAJGD, AM29NG9CDF, YA29NGARAO*)

## 9 Data Availability

The RNA sequencing data presented in this paper are available in the ArrayExpress database (http://www.ebi.ac.uk/arrayexpress) under the accession number E-MTAB-17130.

