## Supplementary Figures for "Integrative phenotypic and transcriptomic validation of an alveolar-like macrophage model reveals early host–pathogen dynamics during *Aspergillus fumigatus* infection"

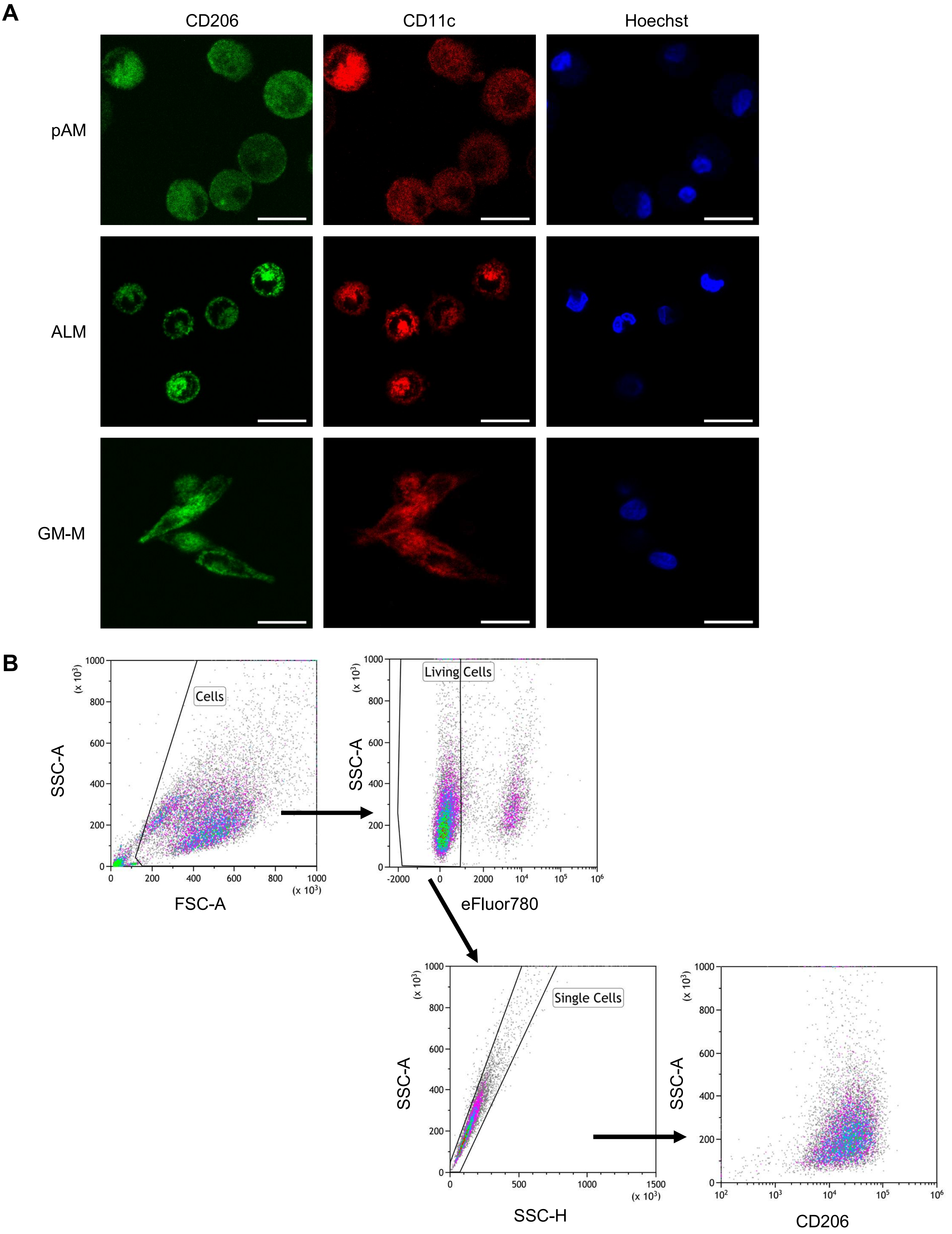


Figure S1: Characterization of the three macrophage populations by immunofluorescence microscopy and flow cytometry.

(A) Immunofluorescence microscopy of pAMs, ALMs and GM-Ms. Single channels of CD206, CD11c, and Hoechst are visualized. Scale bar = 20 µm

(B) Representative gating strategy to examine primary alveolar macrophage markers of pAM, ALM and GM-M.

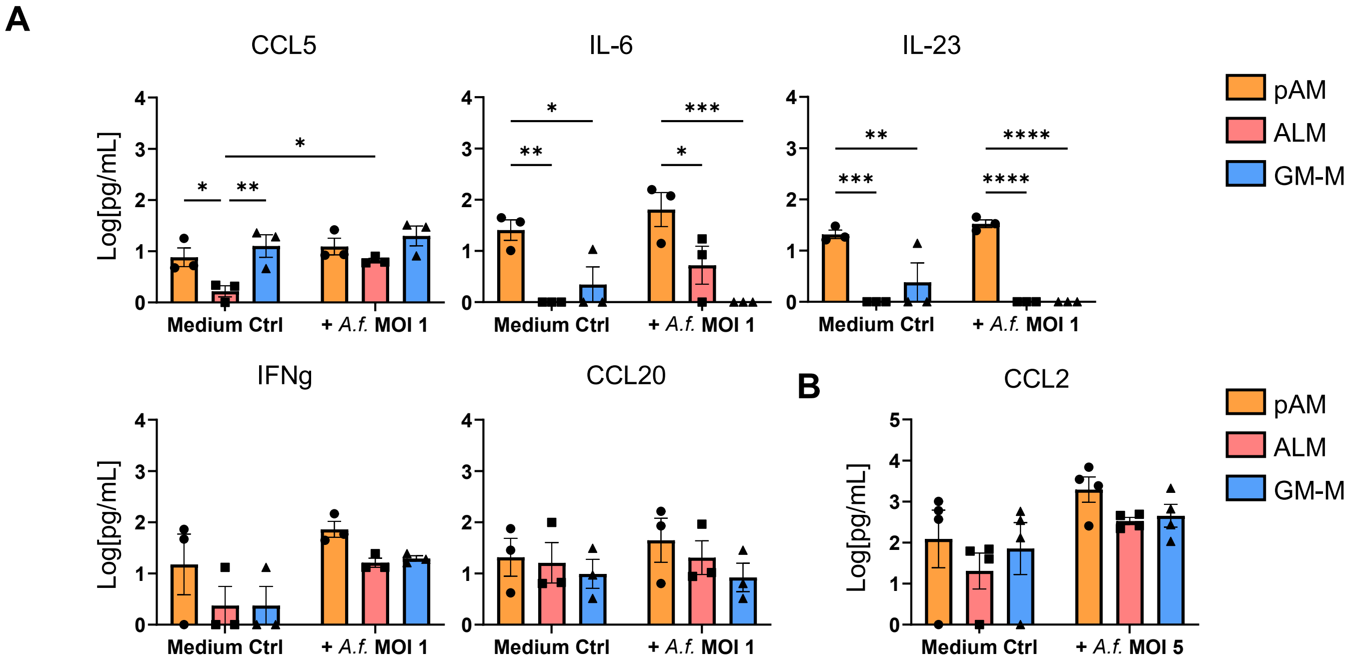


Figure S2: Behavior of pAM, ALM and GM-M during *A. fumigatus* infection.

(A) Quantification of CCL5, IL-6, IL-23, IFNg, and CCL20 secreted by pAM, ALM, and GM-M after 6h of *A. fumigatus* ATCC46645 MOI 1 infection. Data was transformed using log(Y+1). n = 3 +/- SEM, Ordinary One-Way ANOVA, Tukey's Multiple Comparison, * $p < 0.05$, ** $p < 0.01$

(B) CCL2 secreted by pAM, ALM and GM-M during 9h infection of MOI 5 *A. fumigatus* infection. Data was transformed using log(Y+1). n = 4 +/- SEM, Two-Way ANOVA, Tukeys Multiple Comparison; * $p < 0.05$, ** $p < 0.01$, *** $p < 0.001$, **** $p < 0.0001$

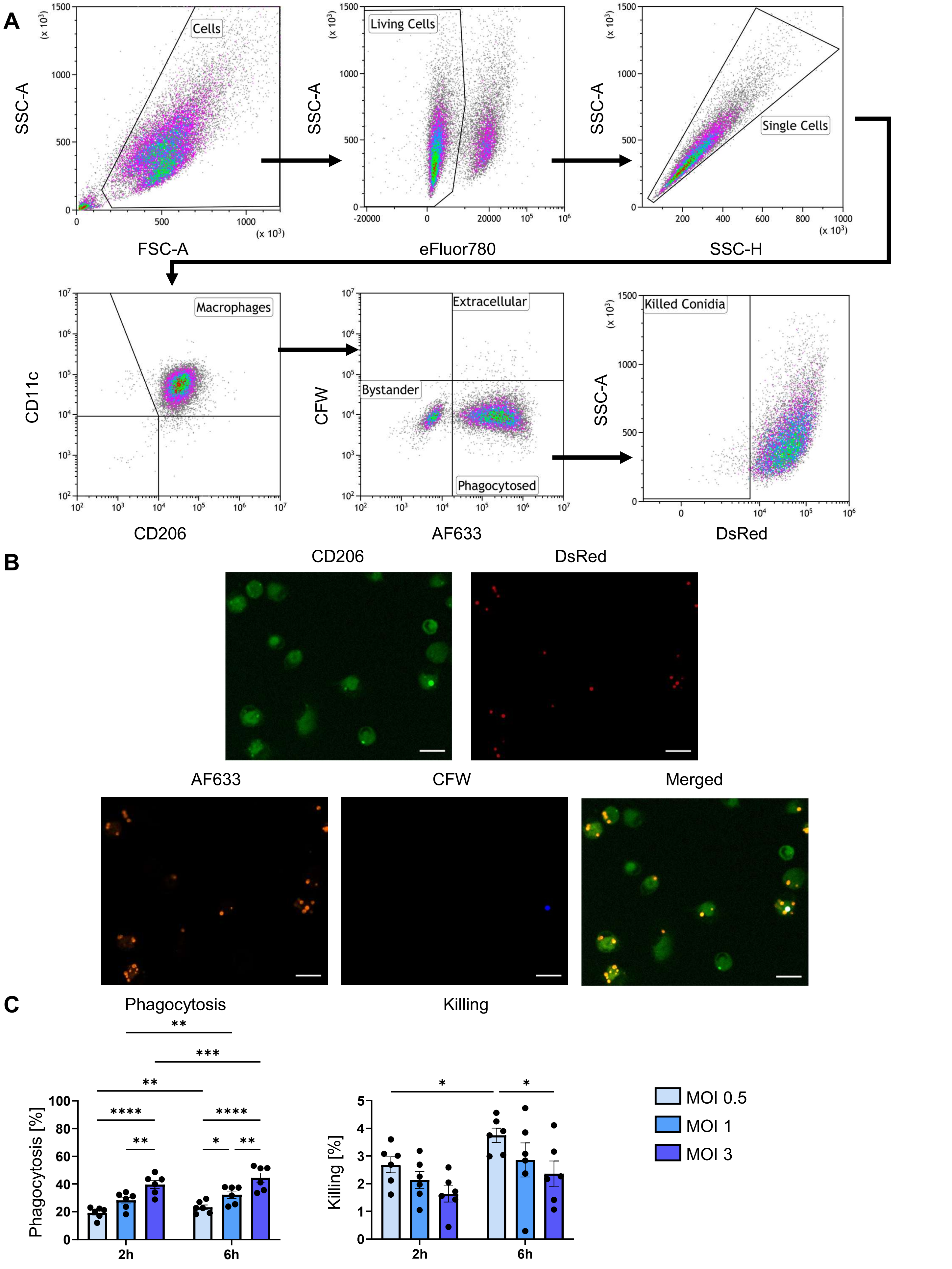


Figure S3: Phagocytic and killing efficacy evaluation using the *A. fumigatus* FLARE strain.

(A) Representative gating strategy to evaluate phagocytosis and killing efficacy of ALM and GM-M infected with *A. fumigatus* FLARE conidia.

(B) Visualization of FLARE infected ALMs via fluorescence microscopy using the ZEISS CLSM780 and a 20x objective. The cells were infected with *A. fumigatus* FLARE MOI 3 for 6h and counterstained with CFW to distinguish between phagocytosed and extracellular conidia. Scale bar = 20 µm

(C) Time dependent differences in phagocytic and killing efficacy of GM-M. n = 6 +/- SEM, Two-Way ANOVA with Tukey's Multiple Comparison, * $p < 0.05$, ** $p < 0.01$, *** $p < 0.001$, **** $p < 0.0001$

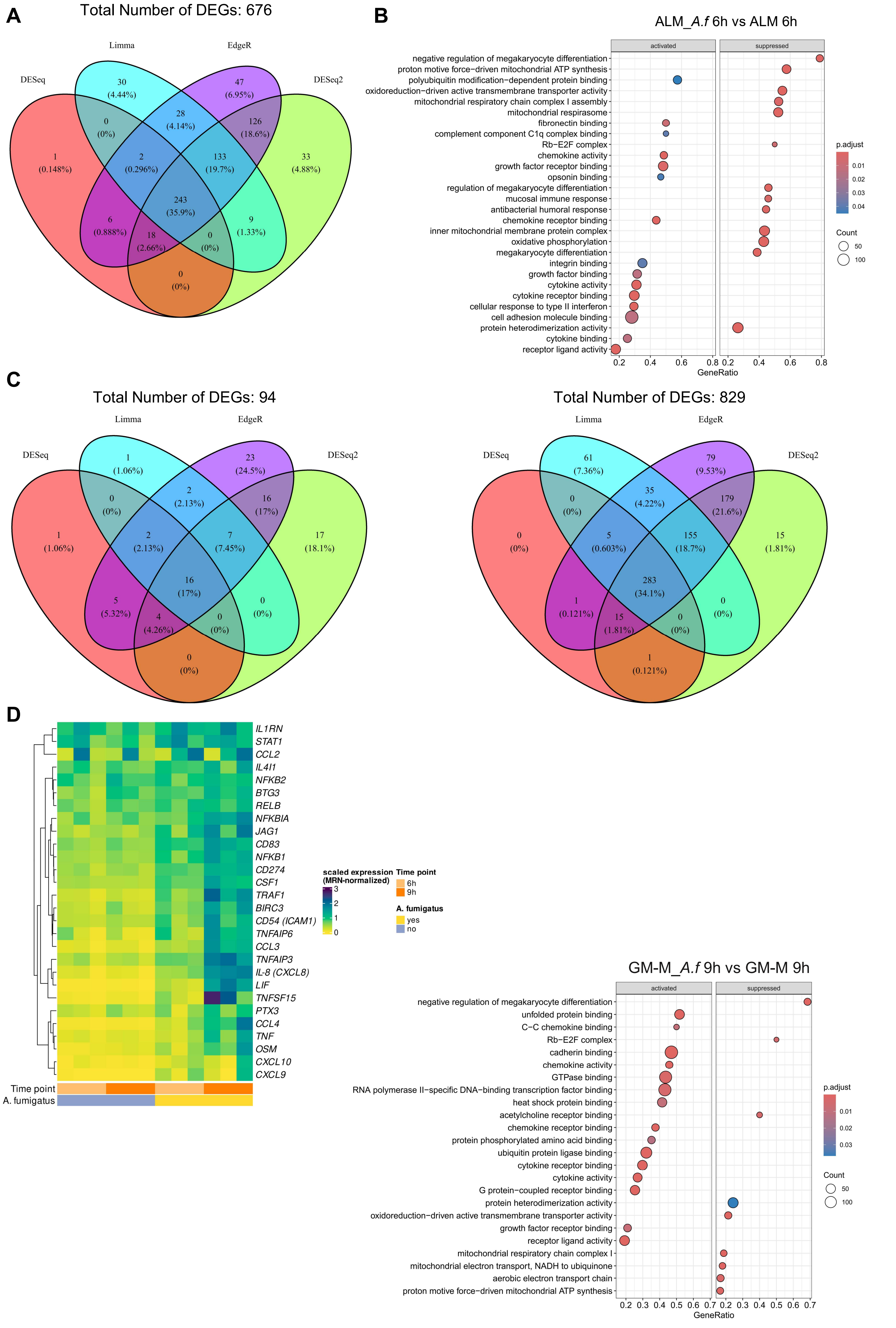
A
Total Number of DEGs: 676
Limma
EdgeR
DESeq
DESeq2
30
(4.44%)
47
(6.95%)
0
(0%)
28
(4.14%)
126
(18.6%)
1
(0.148%)
2
(0.296%)
133
(19.7%)
33
(4.88%)
243
(35.9%)
6
(0.888%)
9
(1.33%)
18
(2.66%)
0
(0%)
0
(0%)
B
ALM_A.f 6h vs ALM 6h
activated
suppressed
negative regulation of megakaryocyte differentiation
proton motive force−driven mitochondrial ATP synthesis
polyubiquitin modification−dependent protein binding
oxidoreduction−driven active transmembrane transporter activity
mitochondrial respiratory chain complex I assembly
mitochondrial respirasome
fibronectin binding
complement component C1q complex binding
Rb−E2F complex
chemokine activity
growth factor receptor binding
opsonin binding
regulation of megakaryocyte differentiation
mucosal immune response
antibacterial humoral response
chemokine receptor binding
inner mitochondrial membrane protein complex
oxidative phosphorylation
megakaryocyte differentiation
integrin binding
growth factor binding
cytokine activity
cytokine receptor binding
cellular response to type II interferon
cell adhesion molecule binding
protein heterodimerization activity
cytokine binding
receptor ligand activity
0.2 0.4 0.6 0.8 0.2 0.4 0.6 0.8
GeneRatio
p.adjust
0.01
0.02
0.03
0.04
Count
50
100
C
Total Number of DEGs: 94
Limma
EdgeR
DESeq
DESeq2
1
(1.06%)
23
(24.5%)
0
(0%)
2
(2.13%)
16
(17%)
1
(1.06%)
2
(2.13%)
7
(7.45%)
17
(18.1%)
16
(17%)
5
(5.32%)
0
(0%)
4
(4.26%)
0
(0%)
0
(0%)
Total Number of DEGs: 829
Limma
EdgeR
DESeq
DESeq2
61
(7.36%)
79
(9.53%)
0
(0%)
35
(4.22%)
179
(21.6%)
0
(0%)
5
(0.603%)
155
(18.7%)
15
(1.81%)
283
(34.1%)
1
(0.121%)
0
(0%)
15
(1.81%)
0
(0%)
1
(0.121%)
D
IL1RN
STAT1
CCL2
IL4I1
NFKB2
BTG3
RELB
NFKBIA
JAG1
CD83
NFKB1
CD274
CSF1
TRAF1
BIRC3
CD54 (ICAM1)
TNFAIP6
CCL3
TNFAIP3
IL-8 (CXCL8)
LIF
TNFSF15
PTX3
CCL4
TNF
OSM
CXCL10
CXCL9
Time point
A. fumigatus
scaled expression
(MRN-normalized)
3
2
1
0
Time point
6h
9h
A. fumigatus
yes
no
GM-M_A.f 9h vs GM-M 9h
activated
suppressed
negative regulation of megakaryocyte differentiation
unfolded protein binding
C−C chemokine binding
Rb−E2F complex
cadherin binding
chemokine activity
GTPase binding
RNA polymerase II−specific DNA−binding transcription factor binding
heat shock protein binding
acetylcholine receptor binding
chemokine receptor binding
protein phosphorylated amino acid binding
ubiquitin protein ligase binding
cytokine receptor binding
cytokine activity
G protein−coupled receptor binding
protein heterodimerization activity
oxidoreduction−driven active transmembrane transporter activity
growth factor receptor binding
receptor ligand activity
mitochondrial respiratory chain complex I
mitochondrial electron transport, NADH to ubiquinone
aerobic electron transport chain
proton motive force−driven mitochondrial ATP synthesis
0.2 0.3 0.4 0.5 0.6 0.7 0.2 0.3 0.4 0.5 0.6 0.7
GeneRatio
p.adjust
0.01
0.02
0.03
Count
50
100

Figure S4: Human transcriptomic profiling reveals fast infection response in ALMs while GM-Ms show attenuated response to *A. fumigatus* infection challenge.

(A) Venn diagram showing the DEGs of ALM infected with *A. fumigatus* ATCC46645 MOI 5 for 6h compared to the uninfected control.

(B) Dot plots visualizing the pathway analysis of ALM infected with *A. fumigatus* ATCC46645 MOI 5 for 6h. The graph displays pathways activated and suppressed by ALM upon fungal challenge.

(C) Venn diagram of DEGs found for GM-M_*A.f* 6h vs GM-M 6h (left) or GM-M_*A.f* 9h vs GM-M 9h (right).

(D) Heatmap displaying the gene expression of GM-M under steady-state or during *A. fumigatus* infection.

(E) Dot plot visualizing the pathway analysis of GM-M infected with *A. fumigatus* ATCC46645 MOI 5 for 9h. The graph displays pathways activated and suppressed by GM-M due to fungal challenge.

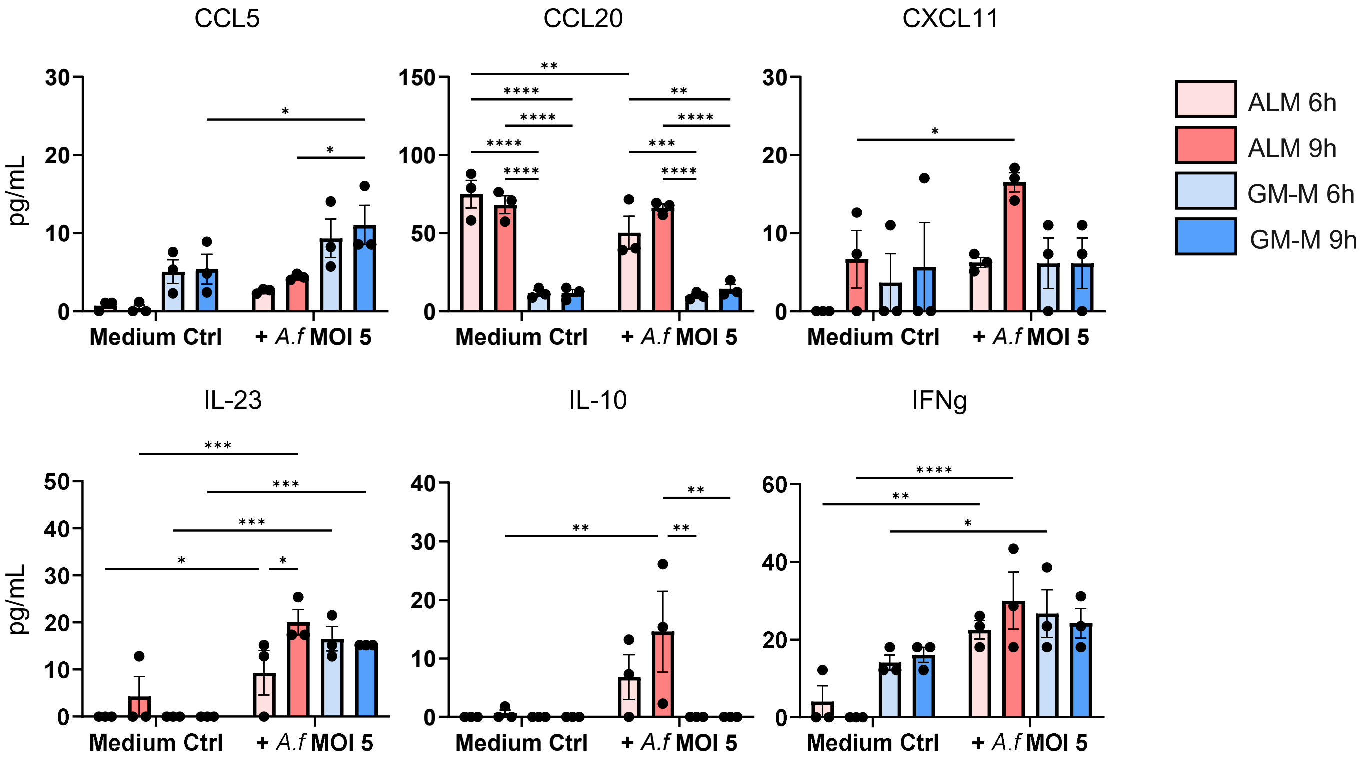


Figure S5: ALM cytokine secretion differs from GM-M. Multiplex analysis of ALM and GM-M donors used during dual RNA-sequencing analysis. ALM and GM-M were left untreated or infected with *A. fumigatus* ATCC46645 MOI 5 for 6h and 9h. n = 3 +/- SEM, Two-Way ANOVA, Tukey's Multiple Comparison; * $p < 0.05$, ** $p < 0.01$, *** $p < 0.001$, **** $p < 0.0001$

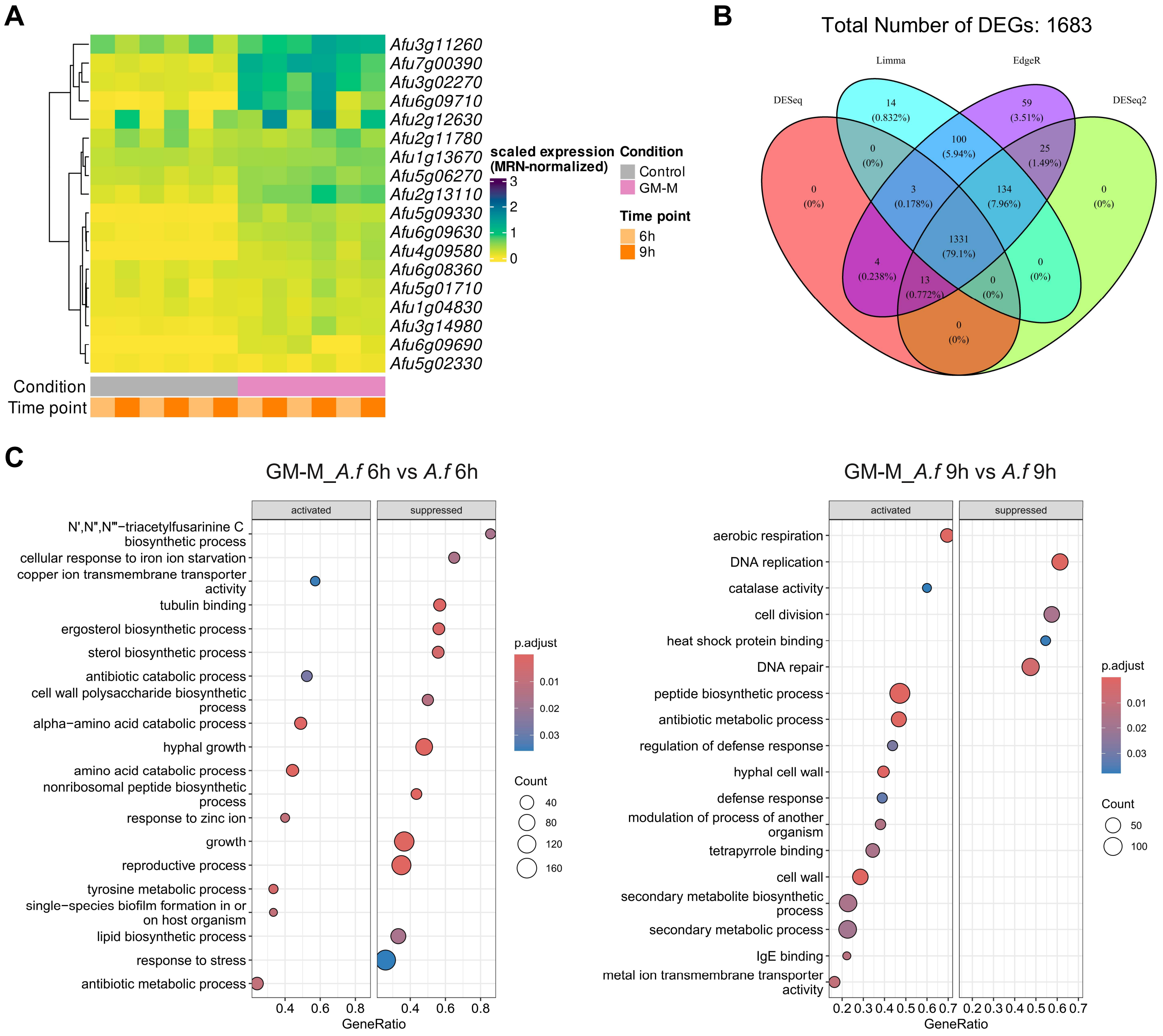


Figure S6: *A. fumigatus* reacts with less prominent counter-defense strategies against GM-M.

(A) Heatmap displaying virulence factors expressed by *A. fumigatus* ATCC46645 due to GM-M challenge as well as the unchallenged control.

(B) Venn diagram of *A. fumigatus* ATCC46645 DEGs after 9h of GM-M challenge.

(C) Representative pathways regulated by *A. fumigatus* after 6h (left) and 9h (right) of GM-M challenge.

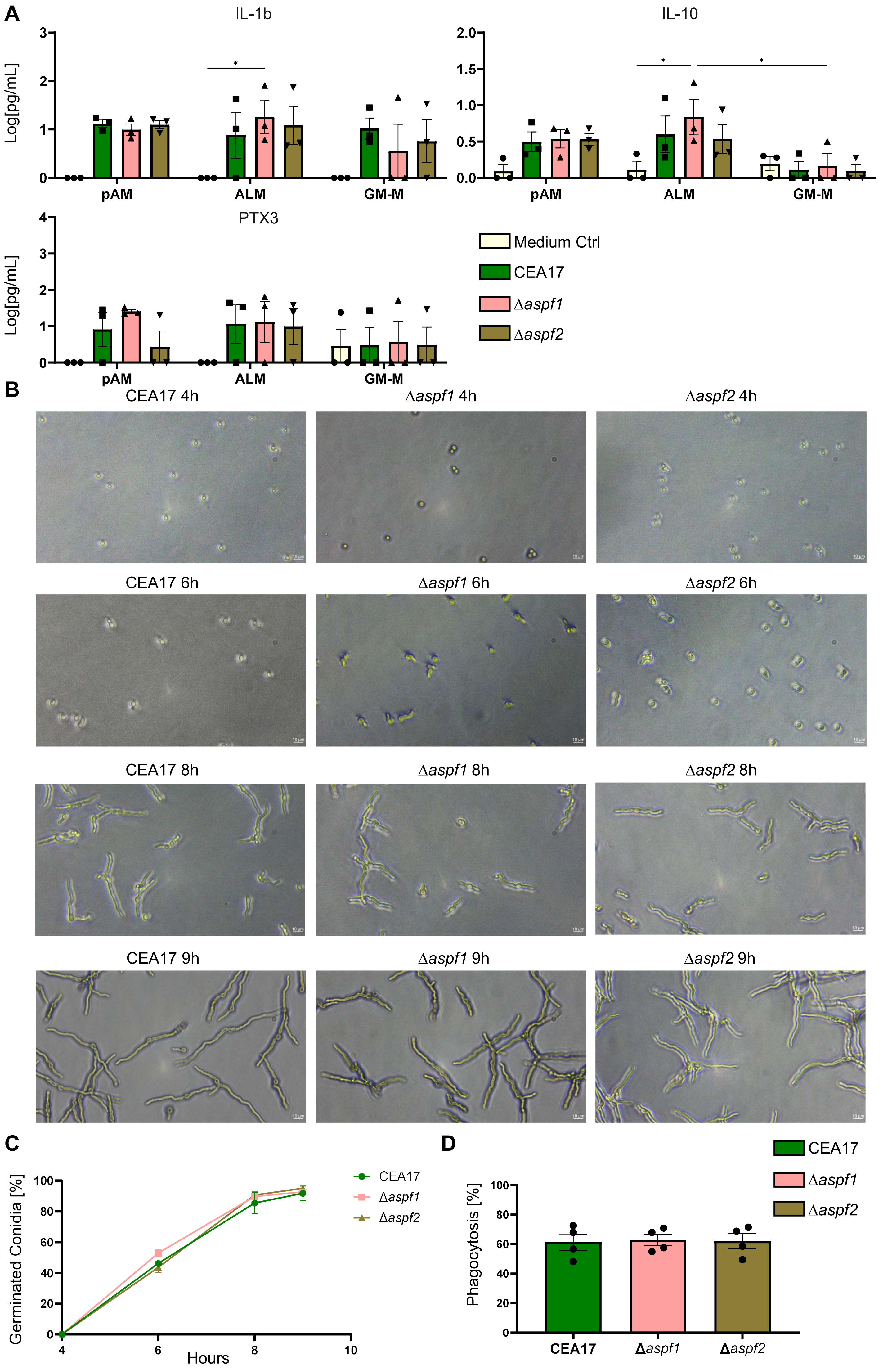
A
IL-1b
Log[pg/mL]
3
2
1
0
*
pAM
ALM
GM-M
IL-10
2.0
1.5
1.0
0.5
0.0
*
*
pAM
ALM
GM-M
PTX3
Log[pg/mL]
4
3
2
1
0
pAM
ALM
GM-M
Medium Ctrl
CEA17
Δaspf1
Δaspf2
B
CEA17 4h
Δaspf1 4h
Δaspf2 4h
CEA17 6h
Δaspf1 6h
Δaspf2 6h
CEA17 8h
Δaspf1 8h
Δaspf2 8h
CEA17 9h
Δaspf1 9h
Δaspf2 9h
C
Germinated Conidia [%]
100
80
60
40
20
0
4
6
8
10
Hours
CEA17
Δaspf1
Δaspf2
D
CEA17
Δaspf1
Δaspf2
Phagocytosis [%]
100
80
60
40
20
0
CEA17
Δaspf1
Δaspf2

Figure S7: *A. fumigatus a*llergens Asp f1 and Asp f2 do not take part in germination and phagocytosis.

(A) CBA analysis of pAM, ALM and GM-M supernatant after 9h of infection with MOI 5 of *A. fumigatus* CEA17, Δ*aspf1* and Δ*aspf2.* n = 3 +/- SEM; Two-Way ANOVA, Tukey's Multiple Comparison; * $p < 0.05$, ** $p < 0.01$, *** $p < 0.001$, **** $p < 0.0001$

(B) Representative pictures of the three *A. fumigatus* CEA17, Δ*aspf1* and Δ*aspf2* strains during germination. Scale bar = 10 µm.

(C) Quantification of germinated conidia after incubation for 4-9h at 37°C, 5% $CO_2$. n = 4 +/- SEM

(D) Phagocytosis of *A. fumigatus* CEA17, Δ*aspf1*, and Δ*aspf2* conidia by ALM after 2h of infection. n = 4 +/- SEM
